# Improved genome inference in the MHC using a population reference graph

**DOI:** 10.1101/006973

**Authors:** Alexander Dilthey, Charles Cox, Zamin Iqbal, Matthew R. Nelson, Gil McVean

## Abstract

In humans and many other species, while much is known about the extent and structure of genetic variation, such information is typically not used in assembling novel genomes. Rather, a single reference is used against which to map reads, which can lead to poor characterisation of regions of high sequence or structural diversity. Here, we introduce a population reference graph, which combines multiple reference sequences as well as catalogues of SNPs and short indels. The genomes of novel samples are reconstructed as paths through the graph using an efficient hidden Markov Model, allowing for recombination between different haplotypes and variants. By applying the method to the 4.5Mb extended MHC region on chromosome 6, combining eight assembled haplotypes, sequences of known classical HLA alleles and 87,640 SNP variants from the 1000 Genomes Project, we demonstrate, using simulations, SNP genotyping, short-read and longread data, how the method improves the accuracy of genome inference. Moreover, the analysis reveals regions where the current set of reference sequences is substantially incomplete, particularly within the Class II region, indicating the need for continued development of reference-quality genome sequences.

---

The current paradigm for analysing human genomes using high throughput sequence (HTS) data is to map to a single haploid reference sequence in which there is no representation of variation^1–3^. Across much of the genome, such exclusion has little effect on the accuracy of genome inference because of the relatively low genetic diversity of humans. However, for some regions, such as the major histocompatibility complex (MHC) on chromosome 6, which contains the human leukocyte antigen (HLA) genes, there is very substantial sequence and structural variation^4^. Such diversity can result in poor genomic characterisation in individuals who carry sequence that is either missing or highly divergent from the single reference. Other locations of high diversity include the KIR^5^ region, olfactory gene clusters^6^, ancient inversions such as that on 17q21.31^7–9^ and regions of recurrent genomic rearrangement^10^, many of which have substantial influence on phenotype and disease risk. In many of these cases, multiple alternative haplotypes have been characterised and are available. For example, there are eight alternative MHC haplotypes in the human reference (GRCh37^11^). More generally, sequencing projects have greatly advanced our understanding of human genetic variation^12–14^; using such information to help characterise human genomes represents an important and unsolved problem.

The problem of the single reference approach and the potential of using known variation to characterise the MHC is demonstrated in Figure 1 for a single individual (CS1). When the standard reference (carrying the PGF MHC haplotype) is used for mapping, large fluctuations in coverage and substantial read mismatches are observed (Fig. 1a, b). However, when a more appropriate reference is used, (here identified by comparing the classical HLA genotypes of the sample with those of the eight reference haplotypes and noting that one of the eight haplotypes was a close match), read coverage and alignment is greatly improved (Fig. 1c, d).

**Figure 1.**
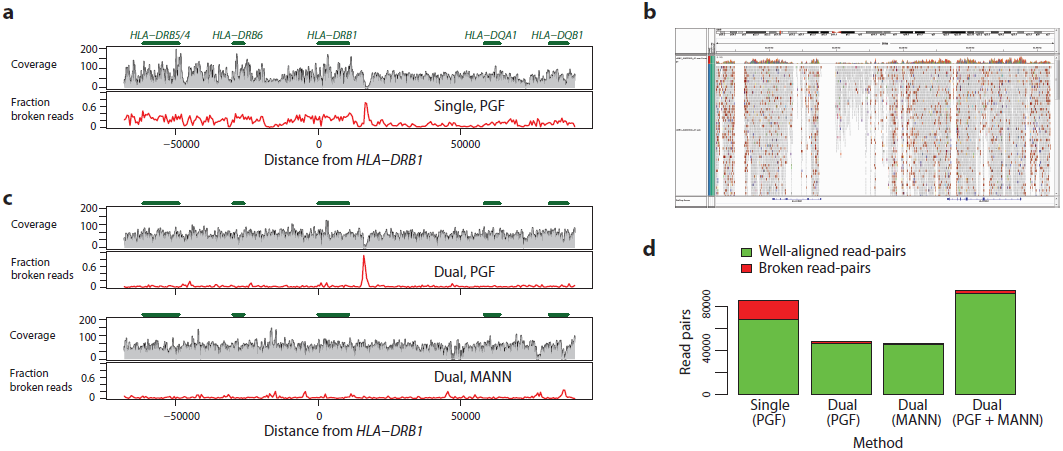
Read-mapping in the MHC Class II region. **a**. Summary of read alignment to a single reference (PGF) for a single sample (CS1) in the MHC Class II region (around *HLA-DRB1*) showing coverage (grey profile) and the proportion of ‘broken’ read-pairs (red line; defined as mapping to different chromosomes; incompatible strands; or implausible insert size). **b** IGV screenshot showing variable coverage and high rate of sequence mismatch for reads aligned in the *HLA-DRB6* / *HLA-DRB1* region. **c** The same metrics as for part a, where mapping has been performed to GRCh37 (including the PGF haplotype) augmented with the MANN haplotype, chosen because the combined classical HLA genotypes from PGF and MANN match those of the sample. **d.** Number of mapped intact (green) and broken (red) read pairs demonstrating that the augmented reference results in many more well-mapped and many fewer broken read-pairs.

In order to use prior information about variation there are five main challenges. First, there needs to be a data structure for representing genomic variation that can accommodate multiple sources of information, from high quality assembled reference sequence (such as the ALT paths in GRCh37) to the lower-quality, but very extensive, catalogues of variation such as the 1000 Genomes Project^12,14^. Second, there have to be algorithms that match high-throughput sequencing (HTS) data from a subsequent sample to the reference structure in order to best characterise the haplotypes present. Third, and potentially simultaneously with step two, there have to be methods for detecting additional variation not yet represented in the reference data structure. Fourth, because most functional information (such as the location and structure of genes) uses the coordinates of a single primary reference, there has to be a coherent way of projecting information from a variation-aware reference onto a primary sequence. Finally, there need to be methods to validate and compare the output from a variation aware reference tool-chain to the genomic information provided by existing approaches that rely on a single reference genome.

To date, limited progress has been made on addressing these challenges. Graph structures of local sequence variation and/or prior knowledge of small-scale variants are used to aid read assembly by several variant-calling algorithms^15–17^. However these make no attempt to build a re-usable reference structure and do not incorporate long or diverged haplotypes such as in the MHC. The pragmatic approach of appending alternative sequences to the end of a reference has been proposed^18^ but cannot solve the fundamental problems of distinguishing between sequence similarity arising through orthology and paralogy. Conversely, the approach of identifying a best reference from among a set has been applied to Arabidopsis^19^, but cannot address the problem that novel genomes are likely to be closest to a mosaic (arising through recombination) of those already known. Methods have been developed to represent multiple aligned genomes in a manner that allows inexact matching^20^, but these are impractical for human genomes because of memory requirements (greater than 1Tb of RAM). Similarly, progress on variation aware data models has been made^21^, though with no implementation. None of these methods represent a general and practical solution to the problem of describing and using information from multiple reference data sets.

Here, we present a solution to the challenges described above. We describe an approach for representing known variation called a population reference graph (PRG) and a series of algorithms that enable characterisation of the genomes present in an individual from HTS data. We build on previous work for using coloured de Bruijn graphs for analysing sequence variation^22^, but also take advantage of the existing tool chain for read mapping and variant calling^3,15^. To demonstrate the value of the method we develop a PRG for the MHC region and combine simulation with analysis of empirical data on SNP genotypes, classical HLA types, short-read and long-read Moleculo data from high coverage samples.

## The population reference graph

A population reference graph (PRG) is a directed acyclic graphical model for genetic variation that is generated by combining information about known allelic relationships between sequences (Figs. 2a, 2b). The graph is constructed in three steps (see Supplementary Text for a detailed description). First, reference sequences are aligned using standard multiple sequence alignment (MSA) methods^23,24^. Second, a graph structure is generated from the MSA by collapsing aligned regions with sequence identity over a defined kmer size. Third, additional SNP information, which is encoded in VCF and related formats through a reference position and alternative sequences, is added to all paths with matching sequence at a given position (for example, a SNP cannot be added to a path where there is a deletion at a given position). Here, all REF and ALT sequences from GRCh37 are used, along with the catalogue of SNPs from the Phase 1 release of the 1000 Genomes Project and the set of classical HLA allele sequences from the International Immunogenetics Information System (IMGT^25^) at key HLA Class I and Class II loci (Table S1). The resulting graph structure can be thought of as a generative model for genome sequences. From a limited set of input sequences, many different paths through the graph are possible, thus mimicking the effect of recombination. In regions of high diversity, the size of the state space can become very large (Fig. S1).

**Figure 2.**
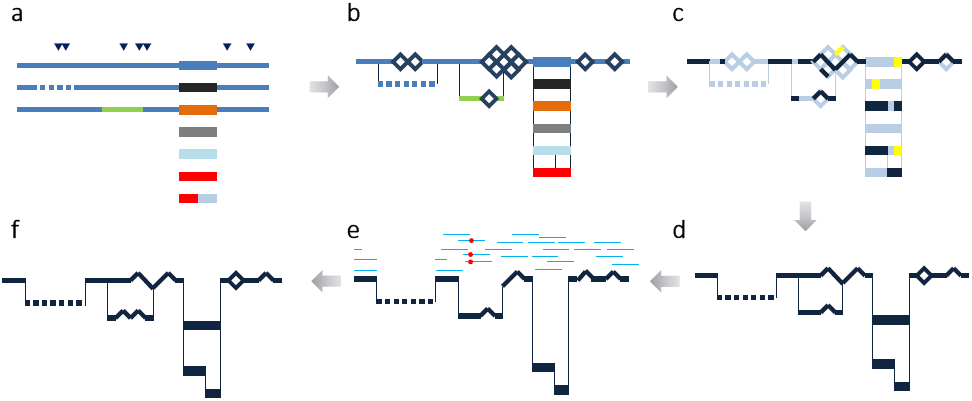
Schematic illustration showing the construction and application of a population reference graph. **a**. Multiple sources of information about genetic variation, including alternative reference haplotypes (lines), classical HLA alleles (rectangles) and SNPs / short indels (triangles) are aligned. Colours indicate divergent sequence, dashes indicate gaps. **b**. A population reference graph (PRG) is constructed from the alignment, resulting in a generative model for variation within the region. SNPs, indicated by diamonds, are added as alternative paths to all valid backgrounds (i.e. excluding sequence with gaps or a third allele at the position). **c**. The PRG is compared to the de Bruijn graph constructed from reads obtained from a sample. Kmers found in the sample are identified (dark blue) along with kmers found elsewhere in the genome that are uninformative about local sequence (yellow). **d**. A hidden Markov model formulation is used to infer the most likely pair of paths through the PRG, allowing for read errors, resulting in an individualised reference chromotype for the sample. **e**. Two haploid genomes are constructed from the reference chromotype and reads (light blue lines) from the sample are aligned, thus identifying places where the sample contains novel variation (red circles; only one path through the chromotype is shown). **f**. Newly-discovered variants modify the reference chromotype, resulting in the inferred chromotype for the sample.

## Using the PRG to infer individual genomes

The development of HTS technologies in humans has largely relied on the notion that the genome(s) of the sample(s) in the experiment will be closely related to those of the reference, thus enabling reads to be mapped accurately and with appropriate certainty. We extend this idea by attempting to infer the (diploid) path through the PRG that most closely resembles the two haplotypes of the sample. Specifically, by comparing the HTS data from a sample to the PRG we construct a diploid personalised reference genome, here referred to as a chromotype. To infer novel variation not yet present in the PRG, we map reads to the chromotype and use existing variant calling software^15^.

Chromotypes are inferred by considering the HTS data from a diploid sample to be emitted by a pair of paths through the PRG and approximating the emission process so as to benefit from the computational efficiency of hidden Markov model techniques. Briefly, HTS data is summarised by the counts of each string of length k (kmer). Similarly, the set of kmers that can be emitted from the PRG is enumerated, eliminating those that occur multiple times across the genome and that are therefore uninformative for local sequence inference (Fig. 2c). Finally, by using a probabilistic model for the emission of kmers (see Methods), the Viterbi-algorithm infers the maximum-likelihood (ML) pair of paths (chromotype) through the PRG (Fig. 2d). Note that this approach does not preserve any phase information across regions where paths merge. The diploid path is thus best understood as a bifurcating/merging sub-graph of the PRG, where heterozygous sites induce bubbles.

To detect novel variation within the sample the inferred chromotype is decomposed into two sequences (with no attempt to establish phase between adjacent bubbles in the chromotype), which are used to replace the homologous region in the primary reference. Reads are mapped to the two resulting reference genomes and each read is placed at its best position across the two reference genomes, as measured by mapping quality. A standard variant caller^15^ is used to discover new alleles independently in the two mappings and a heuristic algorithm modifies the chromotype accordingly. We have also developed an approach for mapping reads directly to the chromotype, which is important for validation using long-read sequences (see below) though currently too slow for primary analysis at the scale of millions of reads.

## Validation and comparison to other methods

To assess the value of the PRG approach in characterising variation within samples we used simulations and empirical data analysis. We compare four approaches to characterising variation.

1. As a base-line we use a single reference (the PGF haplotype within the MHC region from GRCh37) and look at the effect of calling a sample as everywhere homozygous-reference.
2. We use a read-mapping approach (Stampy^3^ followed by Platypus^15^) in which the components were designed explicitly for high sensitivity detection and genotyping of short INDELs and clustered variants.
3. From the PRG, we assess the Viterbi chromotype.
4. From the PRG, we assess the results from mapping reads back to the Viterbi chromotype with BWA^1^ and calling using Platypus^15^ (i.e. the modified chromotype).

The output of each approach can be represented as a chromotype, thus enabling comparison between PRG and single-reference based methods.

### Simulations

We simulated high coverage HTS data (85bp error-free paired-end reads from a 30x genome) for 20 individuals. The primary effect of read errors is to reduce kmer coverage, hence their omission. Each simulated diploid genome consists of two random paths through the PRG for the extended MHC (xMHC). The simulated genomes carry a mixture of recombination events between the original eight MHC haplotypes, SNPs and structural variants of varying size (insertions and deletions from 1 – 125,000bp). We assess the accuracy of the PRG approach through genotype concordance of the inferred paths through the PRG with the simulated paths (Table S2). Across all positions, 99.86% of alleles are correctly recovered. Accuracy at heterozygous SNP positions is similar (99.83%) and drops slightly for INDEL positions (ranging from 95.76% to 99.97%, Figs. 3a, 3b).

**Figure 3:**
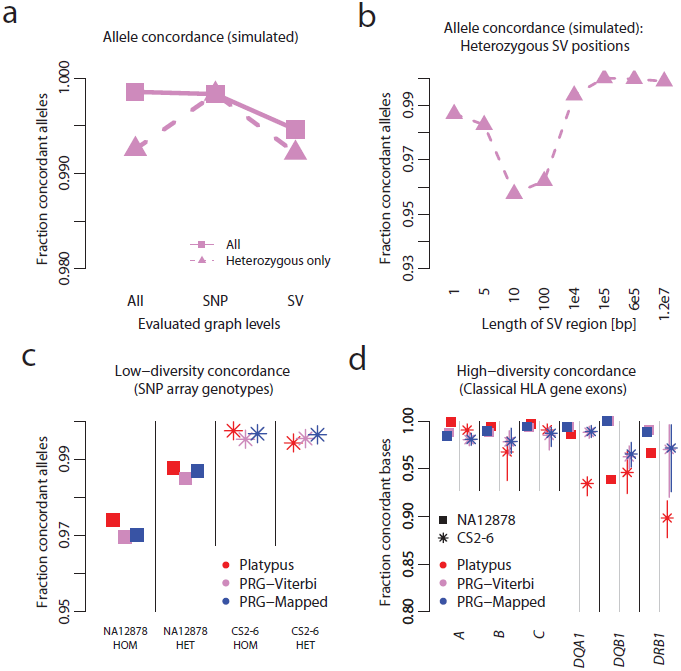
Simulation study and empirical validation. **a.** Concordance between simulated data (20 simulated diploid individuals; 85 bp error-free reads at 30x diploid coverage) and Viterbi path through the PRG stratified by simulated variant type (SNP or indel) and genotype. **b.** Genotype concordance in simulations at sites simulated to be heterozygous for structural variants of different lengths. **c.** Concordance between SNP array genotypes and chromotypes from each method for NA12878 (squares; Illumina Omni 2.5M array) and the CS2-6 samples (stars; Illumina 1M array), stratified by whether the array specifies the genotype as homozygous or heterozygous. Results shown for the mapping-based approach (Platypus, red), the Viterbi-path through the PRG (PRG-Viterbi, pink) and after mapping to the reference chromotype (PRG-Mapped, blue). **d.** Concordance between classical HLA genotypes at *HLA-A*, *-B*, *-C*, *-DQA1*, *-DQB1* and *-DRB1* (measured at a per-base level) and chromotypes from each method for NA12878 and the CS2-6 samples (range of accuracy across CS2-6 displayed as vertical bars). Classical HLA genotypes were generated using sequence-based HLA typing (see Methods).

### Experiment 1: Comparison to SNP array data

To assess the ability of the PRG approach to genotype variation at sites of high uniqueness within the genome, we compared metrics of accuracy at SNP positions independently interrogated through array genotyping and HTS (1 sample [NA12878] at 60x coverage with 100bp paired end-reads and 5 clinical samples [CS2-6] at 30x coverage with 90bp paired-end reads; see Methods).

The accuracy of all approaches is high (Fig. 3a), ≥97.38% concordance with the Illumina Omni 2.5M array (NA12878) and ≥99.53% allele concordance with the Illumina 1M array (CS2-6) for all approaches. Comparing the array genotype concordance of Platypus-generated genotypes and PRG-generated genotypes, we find that both approaches yield comparable accuracies (97.75% vs 97.45% for the 2.5M array and 99.57% vs 99.66% for the 1M array, Fig. 3c, Table S3).

Of the 285 sites at which the array genotypes for NA12878 disagree with the Viterbi chromotype, in 55 cases this difference is driven by the Viterbi chromotype specifying at least one gap character suggesting the presence of an indel that could interfere with array genotyping. We have manually inspected the reference alignment of NA12878 reads for these sites, and find clear evidence for the presence of at least one deletion in 33 of the 55 cases (we provide visualisations of read mapping at all positions in Supplementary File 1). These findings suggest that a significant fraction of the discrepancy between array and PRG approaches results from array errors at polymorphic indels.

### Experiment 2: Comparison to classical HLA data

In regions of high sequence diversity, such as the classical HLA alleles, single-reference mapping and variant calling methods may perform poorly because of the density of mismatches to the reference. To assess the accuracy of different methods at the classical HLA alleles, we compared the per-base diploid genotypes inferred by mapping and PRG approaches to those expected from the results of sequence-based typing (SBT) of the highly polymorphic exons of Class I (*HLA-A*, *-B* and *–C*) and Class II (*HLA-DQA1*, *-DQB1* and *–DRB1*) genes in NA12878 and CS2-6. We analysed agreement with the corresponding allele reference sequence for the reported allele (in HLA nomenclature this means XX:XX:01 or XX:XX:01:01 at 6 or 8 digit resolution respectively)

Where diversity is relatively low and sequence coverage is very high, the accuracy of mapping-based approaches is high; for example 100% concordance for NA12878 at exon 2 of *HLA-A* and *–C* (Fig. 3d, Table S4). However, when coverage is lower, or sequence divergence and / or complications from paralogy are greater (e.g. for *HLA-B* in the CS2-6 samples or *HLA-DRB1* in all samples), the PRG approach typically outperforms mapping (e.g. 97.19% concordance with the PRG genotypes versus 89.85% concordance with mapping-based genotypes at *HLA-DRB1* in the CS2-6 samples). Mapping reads to the chromotype has a marginal effect on accuracy (typically < 1% gain).

### Experiment 3: kmer recovery from high coverage samples

Central to the use of the PRG in assembling individual genomes is the notion that it contains the majority of sequence that is likely to be found in any sample. In the absence of full and independent de novo assemblies, we can assess the extent to which any given chromotype is an accurate representation of the sample by measuring the recovery of kmers from HTS data. We apply this benchmark to NA12878 and the CS2-6 samples.

Across the 4.75 Mb xMHC region, the PGF reference contains 4.52M unique kmers of which 4.8% are not recovered in the HTS data from NA12878 (Fig. 4a). The mapping-based approach predicts 4.94M kmers, of which 1.2% are not recovered while the two PRG approaches predict 4.98M and 4.97M kmers respectively and 0.63% and 0.57% are not recovered. Results are comparable though slightly lower for all methods in the CS2-6 samples (Table S4). Consequently, the PRG approach both predicts greater sequence diversity than the mapping approach and achieves a higher rate of sequence recovery. Although the majority of the xMHC is accessible to all methods, there is substantial spatial heterogeneity in the rate of kmer recovery by all methods (Fig. 4b). Particularly in the HLA class II region, the PRG approach outperforms mapping approaches (Fig. 5), consistent with knowledge of genomic complexity involving the *HLA-DRB* paralogues. We also note that in some regions, in particular distal to *HLA-DRB5*, all approaches perform poorly in terms of kmer recovery (Fig. 5), suggesting that current catalogues of sequence within the xMHC are substantially incomplete.

**Figure 4.**
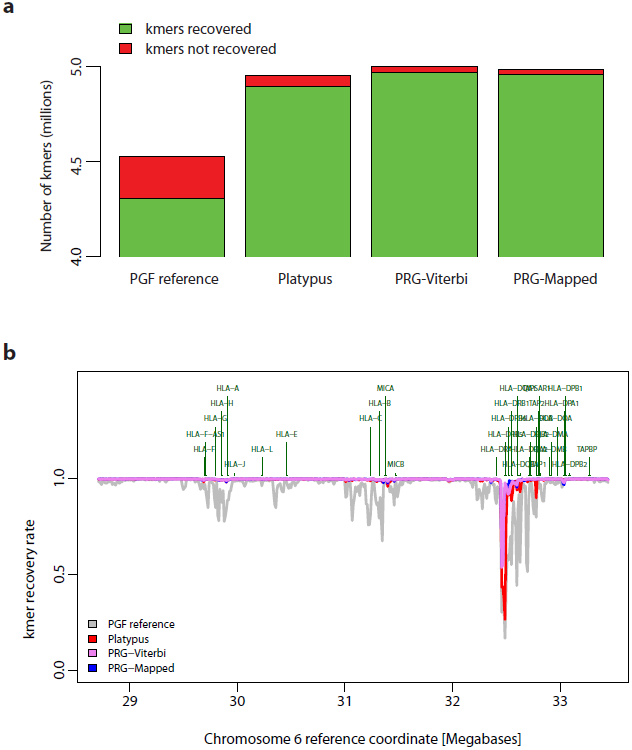
Recovery of chromotype kmers from high throughput sequencing data. **a.** Number of recovered and non-recovered kmers present in chromotypes inferred by the four methods (as for Fig. 3c with addition of single reference represented by the PGF MHC haplotype). A kmer is counted as recovered if it appears in HTS data from NA12878 (∼60x coverage of 100 bp paired-end reads represented by an un-cleaned Cortex^22^ graph; k = 31). Chromotypes are disentangled using a greedy algorithm prior to evaluation, optimizing for the disentangled haplotypes to contain as many kmers recovered in the sample as possible (see Supplementary Text). **b**. Spatial pattern of kmer recovery along the extended MHC region for each of the four chromotypes showing the location of classical HLA loci. Recovery fraction averaged over 1 kb windows.

**Figure 5.**
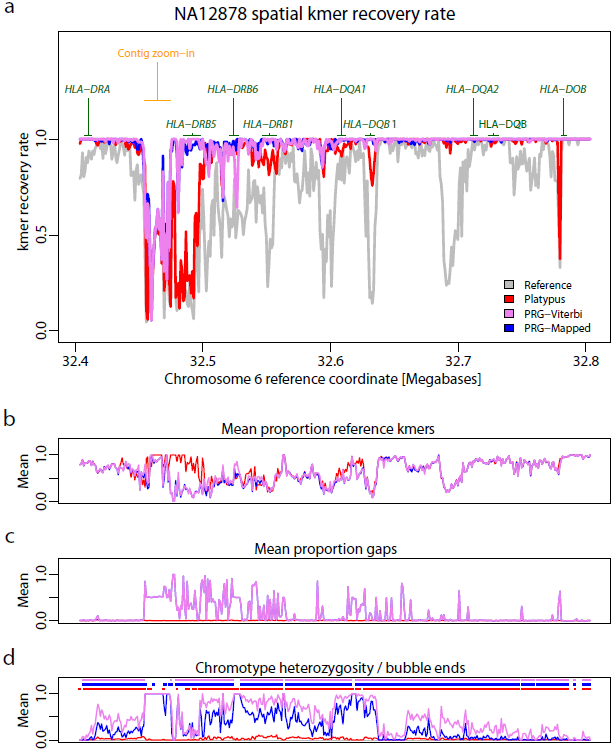
Spatial recovery of kmers within the HLA Class II region. **a.** Blow-up of kmer recovery in Fig 4b in the MHC Class II region for the chromotypes inferred by the four approaches. **b.** Fraction of kmers predicted to be present along region that are also presented in the PGF reference haplotype (1 kb windows; PGF reference not shown). **c.** Fraction of positions in chromotype that correspond to gaps in the multiple sequence alignment used to construct the PRG (1 kb windows). Note that PRG-Complete chromotype is effectively identical to the PRG-Viterbi path. **d.** Fraction of positions in inferred chromotypes that are heterozygous (lines; note this includes sites where one allele is a gap character) and the ending points of chromotype bubbles (points).

Within the classical HLA loci, all methods perform well for class I loci, recovering 98-100% of kmers compared to 80-95% from the PGF reference haplotype (Fig. S2). At Class II loci, however, the advantage of the PRG method is pronounced, with approximately 99% of all kmers recovered for *HLA-DQA1*, *-DQB1* and *-DRB1*, compared to 88-95% for Platypus (against a base-line of 37-85% for the PGF reference haplotype).

### Experiment 4: Comparison to long-read Moleculo data

To assess alternative strategies for genome assembly over longer physical distances, we analysed high coverage long-read Moleculo data (25x coverage) from NA12878 (see Methods). We first identified 29,429 reads (median read length 3,165 bp) likely to have arisen from the MHC region through the presence of diagnostic kmers (see Supplementary Text), then aligned reads to the chromotypes generated by each approach. Read-to-chromotype alignment was performed with a Needleman-Wunsch-like alignment algorithm that aligns to graphs instead of sequence, implemented using dynamic programming (see Supplementary Text). We measure the scaled edit distance between reads and the chromotype (measured as non-identical characters in read to chromotype global alignment, divided by read length in kmers) as an indicator of genome accuracy.

We find that the mapping-based approach achieves the highest number of read alignments with zero mismatches (11,338 vs 10,071 for the modified PRG method). However, both PRG approaches result in significantly fewer reads with many mismatches and/or gaps (Fig. 6a, Table S5). For example, the total number of alignment columns indicating a deletion in the chromotype decreases from 1,017,231 (mapping-based) to 586,852 (modified PRG chromotypes). Likewise, the number of reads with very bad alignments (more than 150,000 gaps in the aligned read or ≥33% of the aligned chromotype string consisting of novel gaps) decreases from 303 to 134. The modified PRG chromotype has a modest benefit over the Viterbi chromotype, increasing the number of perfectly mapped reads from 8,359 to 10,071. Across the *DRB5* region (identified from the kmer recovery analysis as being most poorly represented by the PRG) we find reads that suggest the presence of an inversion relative to known sequence (Fig. 6b).

**Figure 6.**
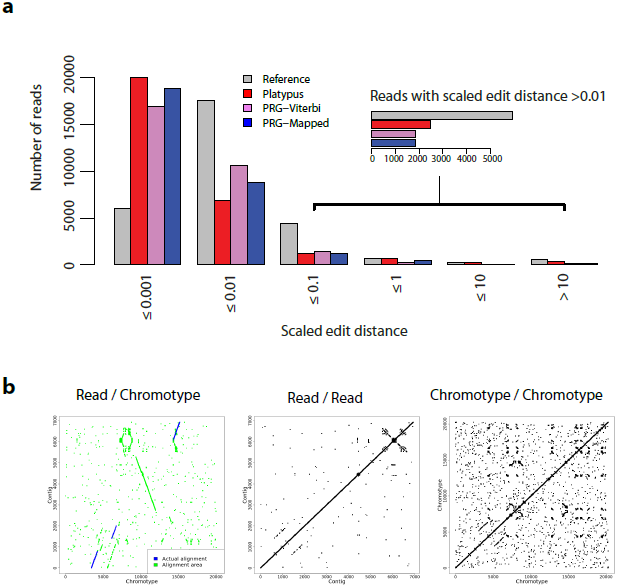
Alignment of long-read data to chromotypes. **a.** Histogram of scaled edit distance (the number of non-concordant columns in the alignment between read and chromotype, divided by the total number of bases in the read) between long-read data (Illumina NA12878 Moleculo xMHC-specific reads, see Supplementary Text) to chromotypes inferred by four methods. Lower boundary for each interval omitted for clarity. Inset shows a blow-up for contigs with scaled edit distance >0.01. **b.** Dot-plot between the sequence of a Moleculo contig and the sequence of the non-gap branch of the Viterbi chromotype for NA12878 over the region highlighted in Fig. 5a (“target region”). There is a point (x, y) if and only if the 10-mer beginning at position x in the chromotype segment is identical to the 10-mer (or its reverse complement) beginning at position y in the Moleculo read. Green indicates the region of the contig which, according to the alignment, is matched to the target region (i.e. each green point represents a contig kmer between the leftmost and the rightmost contig kmers aligned to the target region). Blue indicates that the match between the kmer found at positions x in the chromotype and y in the Moleculo can be recovered from the alignment. Middle, right: Analogous dot-plots for the contig and the chromotype against themselves, showing that there is no large-scale self-similarity along either sequence.

## Discussion

We have presented several innovations that address the problem of how best to represent and use information about known genetic variation in the assembly of HTS data from novel samples. These are an approach for representing information through the population reference graph (PRG), a practical method for reconstructing personalised reference genomes by comparing HTS data to the PRG, a method for detecting variation not present in the PRG and a series of benchmarking tests that enable comparison of methods, whether PRG-based or otherwise, which can complement other benchmarking approaches^26^. Our approach is both modular (such that progress can be made on each element independently) and maintains the ability to project information onto a canonical primary reference that is used as the basis of most functional annotation.

In constructing the PRG, our approach has been to combine multiple sources of reference variation information including GRC37, the 1000 Genomes Project^14^ and IMGT^25^. Importantly, we make no claim that the information contained within the graph is a full or accurate description of the variation within the region. For example, the 1000 Genomes Project variant list is estimated to have a false discovery rate of 1-5% depending on the variant type. However, the presence of false variants within the PRG should have relatively little effect on accuracy, because paths through such variants will have no support in HTS data. The aim of the PRG approach is not to describe all human variation but simply improve genome assembly through representing the diversity of sequence that may be present in an individual. Importantly, we evaluate the approach using metrics that are independent of whether a PRG or simple reference approach is used (kmer recovery, genotype accuracy at known SNP and HLA variants and long-read alignment). Future versions of the PRG could, however, more closely match observed patterns of variation, for example by removing paths never observed and weighting paths by population frequency. However, whether such changes improve genome inference would have to be evaluated empirically. What the approach has demonstrated is that graph based methods offer benefits in regions of high sequence diversity and that there are substantial stretches in the Class II HLA region of the MHC that are poorly characterised by all existing reference material, indicating the need for ongoing collection of reference variation data.

The current implementation has several limitations. First, by summarising information as kmers, we lose longer range information, particularly from read pairs. Likewise, the process of separately mapping reads to each of the pair of chromosomes within the inferred reference chromotype is *ad hoc* and inefficient. Both features arise from our attempts to maximise the usability of the PRG approach, notably efficient inference using HMMs and the desire to use as much of the existing tool-chain as possible. Both problems could also be solved potentially by using richer data structures that retain longer range information in both reference variation data and short-read data, for example the FM-index structure of SGA^27^, augmented de Bruijn graphs that retain short-range path information or mapping directly to the graph. Nevertheless, despite such limitations, we have demonstrated that in regions of the MHC with high structural and sequence diversity, the use of the current implementation can improve genome assembly.

A more fundamental limitation of local graph approaches is that they fail to use the much longer-range information arising from haplotype structure (linkage disequilibrium). Sharing of SNP haplotypes over megabase scales is common even among samples of unrelated individuals, hence such information, which is used in applications such as imputation^28–31^ and refinement of low-coverage sequencing data^32^, has the potential to further improve genome inference. Such information could, however, be included as a prior structure on paths through the PRG, for example in a generalisation of the HMM approaches used by imputation methods^28–31^.

## Methods

A full description of the PRG algorithms can be found in the Supplementary Text, including (i) the algorithms used to build PRGs from a set of reference data, (ii) the algorithmic and statistical methods for inferring a best diploid path (chromotype) through the PRG, (iii) the algorithm to discover novel variation not presented in the PRG, (iv) the graph-mapping algorithm used for the contig analysis. Data used are as follows. CS1 and CS2-6 samples: germ-line DNA was extracted from peripheral blood samples collected from consented clinical trial subjects, previously determined to have evidence of a Class II HLA risk marker for drug induced liver injury^33^. DNA was fragmented and size selected to create 2 × 180 base pair (bp) libraries and 2 × 800 bp libraries. These libraries were sequenced on a HiSeq 2000 to generate 90bp paired end (PE) reads at the BGI (Shenzen, China). For the CS1 sample approximately 200 Gb, and for each of the 5 samples in CS2-6 approximately 100Gb, of sequence was generated. For Fig. 1, CS1 data were initially aligned to GRCh37 on the CLC Genomics Workbench (version 6.5.1) and coverage and intact and broken PE read numbers determined for ∼180 kb surrounding *HLA-DRB1*. This process was repeated utilizing GRCh37 with the addition of the MANN alternative MHC haplotype. For all remaining analyses on CS2-6, reads were mapped to GRCh37 using Stampy^34^ and BWA^1^ and variants were called using Platypus 0.1.8^15^. Read data for NA12878 from the Illumina Platinum genomes project (HiSeq 2000, ∼60x coverage, 100bp paired-end reads; www.illumina.com/platinumgenomes/) was obtained from the EBI (www.ebi.ac.uk/ena/data/view/ERP001775). Reads were aligned to GRCh37 using BWA 0.6.2^1^ and variants were called with Platypus 0.1.8^15^. Moleculo data is available from the 1000 Genomes Project ftp site; ftp.1000genomes.ebi.ac.uk/vol1/ftp/technical/working/20131209_na12878_moleculo.

## Acknowledgements

Funded by grants from GSK and 100956/Z/13/Z from the Wellcome Trust to GM, a Nuffield Department of Medicine Fellowship to ZI, and a Sir Henry Dale Fellowship jointly awarded by the Wellcome Trust and the Royal Society to ZI (102541/Z/13/Z). We thank Mike Eberle and colleagues at Illumina for early access to the Moleculo data.

## Supplementary Methods

In this document, we give precise descriptions of the algorithms for population reference graphs (PRGs).

### High-level summary

We describe the following subsequent steps:

1. Construction of Population Reference Graphs (PRGs) to represent multiple genomes:

- How to create a nucleotide PRG from a catalogue of variation, consisting of scaffold haplotypes for the whole region to be modeled and additional variant specifiers.
2. Genome inference:

- How to create a kMer-PRG, a kMer-emitting object with identical haplotype structure to nucleotide PRG.
- How to compress the kMer-PRG in a way that reduces its size and improves its statistical properties for the model we describe (the compressed kMer-PRG is called “multi-PRG”, because states of the model can now emit multiple kMers).
- How to define and parameterize a Hidden Markov Model (HMM) “on top” of the multi-PRG.
- How to use the HMM to infer a sample’s chromotypes, and how to represent them in VCF format.
3. Novel variant detection:

- How to take pairs of inferred haplotypes (from the HMM on top of the multi-PRG) and use them as a basis for conventional mapping technologies, enabling the discovery of variants not present in the original catalogue of variation.
4. Graph alignment and validation:

- How to align sequence to PRGs, using a modified version of the Needleman-Wunsch algorithm.
5. Experimental details:

- The protocol followed to create a PRG for the extended human MHC.
- Sample data, HLA types and availability.

## 1 Constructing a nucleotide PRG

We define a population reference graph as a directed acyclic graph with one designated start vertex and a set of designated final vertices. There is a *level function* that returns a positive integer for each vertex; all edges are defined between vertices of consecutive levels; and all final vertices are of the same level. All edges are labeled either with a nucleotide or a gap symbol, and that label is emitted upon traversal. Each node has a (potentially improper) probability distribution over the edges emanating from it.

The algorithm for building PRGs is based on a catalogue of scaffold haplotypes and additional variant specifiers (catalogue of variation, COV). Scaffold haplotypes span the region to be modeled, additional variant specifiers define allelic variation in the context of one or more of the scaffold haplotypes. SNPs, for example, are typically included as additional variant specifiers.

Informally, we will construct a PRG that contains all scaffold haplotypes as paths, allowing for recombination between a set of haplotypes whenever there is a stretch of identity between the members of the set. Additional variant specifiers are represented as “bubbles” on top of the scaffold haplotype graph.

PRG definitions and generative algorithm:

- Let the directed connected graph G consist of the set of directed edges E and the set of vertices V, i.e. G = (V, E). For all e = (v_x_, v_y_) ∈ E, v_x_ ∈ V and v_y_ ∈ V; we call e the directed edge from v_x_ to v_y_. For PRGs, we require that there is a well-defined *level function* function l(v) for each node v, according to the following definition. There is one and only one vertex v_0_ with no incoming edges and l(v_0_) = 0. This vertex is called the start vertex. For every (v_x_, v_y_) ∈ E, we define l(v_y_) = l(v_x_) + 1. All vertices with no outgoing edges are called “final vertices”, and we require that the level function returns the same value L for all final vertices.
- Each edge e ∈ E is labeled with a nucleotide (A, C, G, T), a gap symbol (“_”) or possibly a wildcard character (“*”), which stands for any nucleotide. At each node n, there exists a (possibly improper) edge probability distribution over the edges emanating from that node, specifying the probability P_n_(e) to follow edge e, conditional on being at node n. (For notational clarity, we do not explicitly consider the case here that there can be multiple edges between two nodes with different labels, or with the same label. However, all definitions made in this document are easily extended to cover these cases.)
- To generate a haplotype from that model, carry out the following algorithm:

1. Define a “current vertex” variable c_v_ and initialize it as c_v_:= v_0_.
2. Select an edge (v_x_, v_y_) from the set of edges emanating from c_v_ according to the edge probability distribution at c_v_. Emit the label of the edge and set c_v_ = v_y_.
3. If c_v_ is a final vertex, terminate; otherwise, go to step 2.
- The model so-specified is similar to the well-known class of haplotype graph models [1]. In particular, definition of a suitable emission probability distribution on top of each edge label will result in a Hidden Markov Model (HMM); and the model can easily be generalized to emitting diploid data. For details, see [1, 2].

PRG construction algorithm, introductory definitions:

A. Start with a multiple sequence alignment (MSA) of all S_N_ scaffold haplotypes (constructed using external software, see section “Sequence Alignment”). The MSA has S_N_ rows and L columns (L depends on the exact scoring configuration and algorithm used for creating the MSA; also, L is also the last level of the PRG to be constructed). Each level i > 0 in the PRG refers to the i-th column of the MSA.
B. S_n,i_ denotes the i-th position of the row of the MSA for haplotype n. (n ∈ {1 .. S_N_}, i ∈ {1 .. L}).
C. There is a set X of additional unique variant specifiers of the form (n,i_1_,i_2_,*seq*). n specifies which row of the MSA (which scaffold haplotype) the variant specifier refers to; i_1_ and i_2_ define the column range within the MSA that the variant specifier refers to; and *seq* defines an alternative sequence of characters (constrained to the set of nucleotides, “_” and “*”) to be available at these columns. For example, a SNP could be specified as (2, 5, 5, “A”): this means that position 5 in scaffold haplotype 2 might also be an “A”, instead of, say, a “T”. Although in principle fully generalizable, the current version of the algorithm is specified only for variant specifiers of length 1 (i.e. i_1_ = i_2_); however, it is possible to specify independent variant specifiers at consecutive positions. Overlapping variant specifiers are not permitted.
D. For each vertex v, we define a function H(v) which specifies a set of scaffold haplotypes “attached” to v. At each level of the PRG, each scaffold haplotype is attached to exactly one node. Informally, the set of attached scaffold haplotypes will determine which labels the edges reachable from n will carry. We also define a function *suffix*(v, r) for each vertex v at level l(v) ≤ (L – r) which returns the set of strings defined by

a. For all n ∈ H(v), the concatenated symbols from columns l(v) + 1 .. l(v) + r of row n of the MSA (i.e. from the scaffold haplotypes attached to v).
b. The strings from a), modified by the set of relevant variant specifiers. Each distinct variant identifier for a particular position (i.e. relating to the same scaffold haplotype; at the same position; with a distinct *seq*) maps to at least one additional string, generated by substituting the specified position in the string from a) with the allele specified by the variant specifier. For performance reasons, however, we ignore recombination between variant specifier at different positions (this only applies to the function *suffix*(..), and <u>not</u> to the PRG itself). For example, two nearby tri-allelic positions might induce *suffix(..)* to return three strings. We note that the effect of this simplification is that suffix distributions from different nodes might look more distinct than they really are, but never more similar.

PRG construction algorithm:

1. Construct a multiple sequence alignment from the scaffold haplotypes (using external software, see section “Sequence Alignment”).
2. Initialize the PRG with a single start vertex v_0_ at level 0 and set H(v_0_) to be the set of all scaffold haplotypes.
3. Iterate from i = 1 .. L:

a. For each vertex v at level (i – 1):

- For all scaffold haplotypes with index n from the set H(v): Determine the character S_n,i_ of the corresponding position / haplotype in the MSA and the set {(n,i,i,*seq*)} of relevant additional variant specifiers. If there exists an edge (v, v’) carrying the character S_n,i_, follow that edge and add n to the set H(v’). Otherwise, create a new vertex v’ with l(v’) = i and the corresponding edges (v, v’), labeled with the S_n,i_. Further ensure that for each element {(n,i,i,*seq*)} there is an edge (v, v’) labeled with *seq*.
b. For all possible pairs (v_x_, v_y_) of (newly created) vertices at level i: We merge v_x_ and v_y_ if their suffix distributions *suffix*(v_x_, r) and *suffix*(v_y_, r) are similar. We use r = 20 and require that all r compared positions are non-gaps (i.e. ≠ “_”). If some of the r compared positions are gaps, we dynamically increase r (due to performance only up to a threshold– we assume that two suffix distributions are not identical if we reach this point). “Similar” in the currently implementation means “identical”. If the set of scaffold haplotypes was large and representative of a population, using similarity measures similar to those used when constructing haplotype graphs would be an alternative ([1, 2]). Suffixes that contain wildcard characters are treated in a special way. We define that vertices with suffix distributions that contain only wildcard characters (“*”) are compatible with all other nodes; and that single suffix strings consisting exclusively of wildcard characters are not considered when determining suffix compatibility. To merge two vertices v_x_ and v_y_, define H(v_x_) = H(v_x_) ∪ H(v_y_); redirect all edges going into v_y_ to v_x_ (and remove duplicate edges with identical labels, if necessary); and delete v_y_.
c. If any vertices were merged during step b), repeat step b).
4. We have now constructed a PRG for the complete MSA. For post-processing,

a. Mark all nodes and edges that are necessary to trace the path of the scaffold haplotype which is identical to the canonical reference genome in the region through the graph.
b. Identify and remove arcs (sequences of connected nodes and edges, equivalent to a subpath through the graph; see Section “kMerification” for a formal definition of “subpath”) that consist exclusively of edges labeled with wildcard symbols (“*”) and that can be removed without affecting the reachability of any subpath that does not exclusively consist of wildcard symbols. (We remove as many wildcard symbols as possible because we ignore them in downstream analyses).

### 1.1 Varying number of scaffold haplotypes

The algorithm specified assumes a fixed number of scaffold haplotypes across the region to be modeled. This is, however, not always the case: in the extended MHC, for example, we have eight scaffold haplotypes across the region (see Section “Graph for the extended MHC”), but a much higher number of haplotypes for the six classical HLA loci.

We deal with such situations by dividing the whole region to be modeled in stretches with an identical number of scaffold haplotypes; we then construct separate PRGs for each stretch; and finally we connect the separate PRGs (by fusing last-level nodes of one PRG with the first-level nodes of the next PRG) to obtain a combined region-wide PRG.

## 2 Genome inference

Before giving a formal description of the process of genome inference, we give an informal introduction.

The object we have built – a nucleotide PRG – models population sequence variation at the level of individual nucleotides. While having many advantages (immediately intuitive representation of sequence; easy to visualize; clear preservation of gap homology from the MSA), there are also disadvantages to this approach. Making genome inference (i.e. computing the two most likely paths through the PRG for some given sample data, and detection of additional variation) from the nucleotide PRG itself is not straightforward, for we don't directly observe its most fundamental unit, the nucleotide in its full PRG-level context, from sequencing data.

There are two ways around that:

1. Computation of all possible micro-haplotypes (kMers) of specified length through the nucleotide PRG and counting how often each micro-haplotype occurs in sample data. We refer to this process as “kMerification” and use it as our main approach for genome inference.
2. Read alignment – in analogy to read-to-reference alignment, it is possible to align sequencing reads to a PRG, resulting in a labeling of each individual nucleotide with the level of the PRG it is assumed to be homologous to. Read alignment has many advantages over kMerification – it is more tolerant of sequencing errors and utilizes the full length of sequencing reads to establish homology. However, at least in our current implementations, it is much slower than kMerification-based inference. Hence we use read-to-graph alignment to validate (see Section 4) the genomes we have inferred, but not as a primary means of genome inference.

As stated above, we rely on kMerification as the base for genome inference.

That is, we “kMerify” the nucleotide PRG to obtain a kMer-PRG, a graph with equivalent haplotype structure but edges labeled with kMers instead of individual nucleotides.

To improve some statistical properties of that graph for downstream inference, we further “edge-compress” the kMer-PRG to obtain an object we refer to as “multi-PRG”. The main difference is that we have combined all non-branching stretches of levels in the kMer-PRG into single levels for the multi-PRG, with edges labeled with multiple kMers.

Finally, by assuming that observed kMer counts in the sample data follow a Poisson distribution parameterized by how often a kMer appears in the genome (i.e., after some pruning: how often a kMer appears in a state of the multi-PRG), we use the multi-PRG to derive an efficient Hidden Markov Model (HMM) to infer an individual’s two assumed haplotypes.

The following figure illustrates the different steps of transformation from nucleotide PRG to multi-PRG.

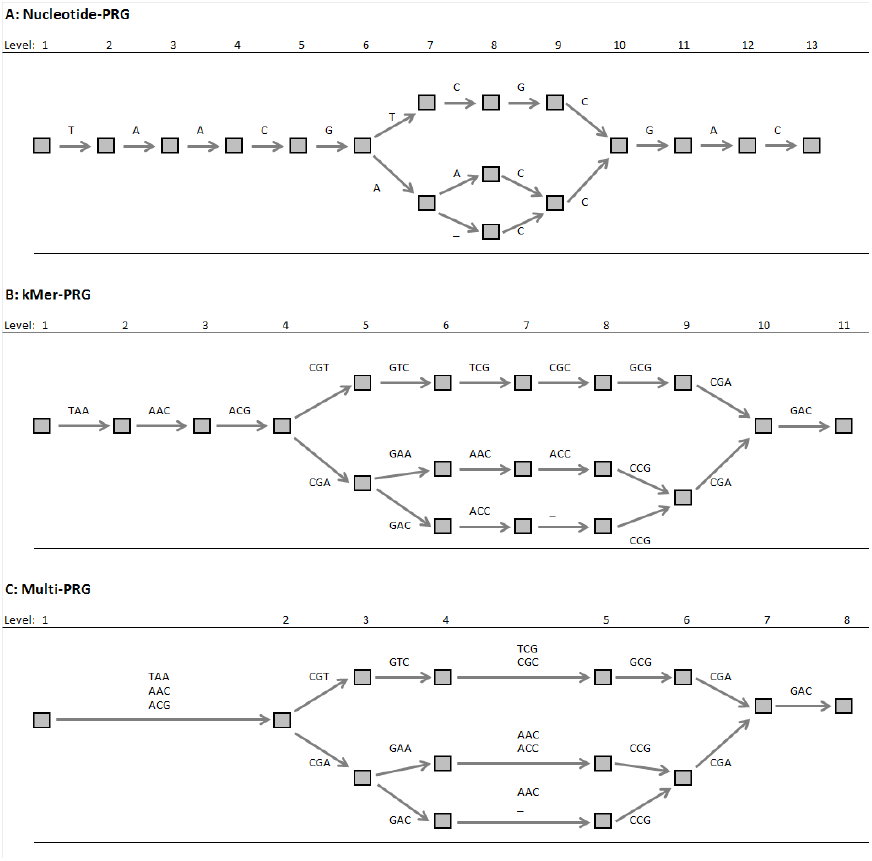

### 2.1 kMerification

The PRG has a probability distribution over the space of possible emitted haplotypes. Each possible haplotype can be transformed into a sequence of kMers. A PRG thus also induces a probability distribution over possible sets of emitted kMers. We now describe the algorithm to transform the PRG into a kMer-PRG that has the same distribution over the set of possible haplotypes and hence over the possible sets of emitted kMers.

Informally, we are searching for an object with equivalent haplotype output distribution, but which explicitly specifies kMers. We call that process “kMerification”.

In order to kMerify a nucleotide PRG, we need to define the set of nodes and edges that the kMer-PRG is to consist of. We define the edges by finding all “micro-haplotypes” of length k (i.e. subwalks of k non-gap characters) specified by the nucleotide PRG. In the process of kMerification, each such micro-haplotype will become one edge in the kMer-PRG.

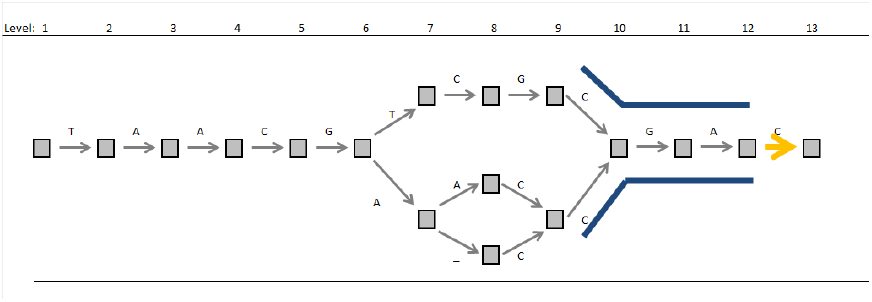

Consider, for example, the two subwalks of length 3 starting at level 9 marked with blue lines in the figure above. In the kMer-PRG, both walks will be represented by separate edges. Note that there will be two edges, even though they will be labeled with the same kMer – this is because the underlying walks through the nucleotide PRG are different.

We now have an intuitive understanding of the edges that will be present in the kMer-PRG, but what about the nodes? Initially assume that (at a given level) each edge in the kMer-PRG ends in its own node. In order to ensure that the haplotype structure of the kMer-PRG reflects that of the nucleotide PRG, the question now becomes: which of these nodes have to be merged? Or, expressed differently: which kMer-PRG edges (reflecting walks of length k) need to end in the same kMer-PRG node?

In order to analyze that question for our two example paths, consider subwalks of length k + 1 from level 9. In the figure above, these walks can be constructed by elongating the two blue walks of length k by adding the orange edge. We note that the two subwalks of length k + 1 are, apart from the first component, identical: by definition the next level of the kMer-PRG will contain exactly one edge to represent the corresponding walk of length k. From the structure of the nucleotide PRG, it is also clear that this one kMer-PRG edge (starting at level 10 in the nucleotide PRG) needs to be connected to the two original kMer-PRG edges (starting at level 9 in the nucleotide PRG). Hence the two original kMer-PRG edges starting at level 9 in the nucleotide PRG need to end in one node, and the kMer-PRG edge starting at level 10 in the nucleotide PRG will start at that node.

Expressed more formally, at each level each edge in the kMer PRG, representing one walk through the nucleotide PRG, ends in a separate node, unless there is second edge in the kMer PRG at the same level which represents a walk that is, apart from the first component, identical to the walk of the first edge.

kMerification algorithm, introductory definitions:

A. A kMer is defined as a word (string) of length k, consisting of the characters A, C, G, T and the symbol for ambiguity (“*”), or as a string of length 0. We refer to 0-length kMers as “gap” kMers. To transform a sequence of characters (consisting of the four nucleotide characters, gap (“_”) and ambiguity (“*”) symbols) of length L into a sequence of kMers, carry out the following algorithm for i = 1 .. (L-k+1): Starting from i and extending to the right, extract a substring of length >= k which contains exactly k non-gap characters. If the first symbol of the so-extracted substring is the “gap” symbol, we define the kMer emanating from position i to be the “gap” kMer. Otherwise, remove all “gap” symbols from the substring and we define the kMer emanating from position i to be equivalent to the substring. (To give an example, transformation of the sequence AC_TAG into kMers of length 3 yields: “ACT”, “CTA”, “_”, “TAG”. “_” represents the gap kMer.) Whenever a kMer contains one or more wildcard characters, we substitute all non-wildcard symbols in the kMer with the wildcard character. (One wildcard character makes the whole kMer ambiguous; we make that definition because it simplifies downstream statistical analyses.)
B. The definition of kMer-PRGs is identical to that of PRGs, with the exception that edges are labeled with kMers of length k (“kMer-edges”). k remains constant across a kMer-PRG.
C. A subpath in the PRG is defined as a sequence of edges e_1_, e_2_ … of the structure e_1_ = (v_x_, v_y_), e_2_ = (v_y_, v_z_) … (we could thus also call a subpath a “walk” through the graph). The set of possible subpaths from the start vertex to one of the final vertices defines the set of possible emitted haplotypes. We could measure the length of a subpath by either the total number of included edges or by the length of included edges which carry non-gap characters. Here we use the second measure (kMerification has to “jump over” gap characters), i.e. we define the length of a subpath to be the number of included edges that are not labeled with gap characters. A subpath defines a sequence of emitted symbols and can thus be transformed into a sequence of kMers (if it is long enough). Specifically, a subpath of length k induces a kMer of length k).
D. It is clear that each subpath of length k in the PRG has to exist as one kMer-edge in the kMer-PRG. While building the kMer-PRG, we keep track of each kMer-edge’s underlying subpath and of the nodes and edges in the PRG that this subpath traverses.
E. We define a function K(v) for all vertices v of the PRG that specifies a set of kMer-edges of the kMer-PRG that are attached to v. A kMer-edge can be attached to a node in the normal PRG if v is its underlying subpath’s second node (but not all kMer-edges have to be attached to a vertex). In implementing the algorithm, K(v) is the empty set for all vertices in the beginning and then extended iteratively while creating the kMer-PRG.

We now describe the core algorithm for kMerification. kMerification works incrementally, i.e. we will move through the nucleotide PRG from left to right, creating nodes and edges of the kMer-PRG as we walk along.

Informally, to define the initial set of kMer edges, we carry out a forward search for subpaths of length k starting at v_0_ of the PRG.

Then, conditional on an existing set of edges of the kMer-PRG (each representing a walk ending in a node in the nucleotide PRG), we create the next level of kMer-PRG edges by extending the walks of the edges of the present level by one character each (or by all possible one-character extensions, if there are multiple possible extensions).

To define the set of nodes required at each level of the kMer-PRG, we remove the first element from the walks (through the nucleotide PRG) corresponding to the kMer-PRGs edges at that level, and we create one kMer-PRG node for each unique reduced walk.

We stop once we have covered the complete PRG.

kMerification algorithm:

1. We initialize the kMer-PRG by creating a start vertex v_0;kMer_ (we use the additional subscript to distinguish between v_0_ of the PRG and the start vertex of the kMer-PRG). We carry out a forward search (using standard algorithms) for subpaths of length k starting at v_0_ of the PRG. We create a kMer-edge for each found subpath, transform the underlying subpath sequence into a kMer and use this kMer as a label for the edge. We attach each created kMer-edge of the form (v_0;kMer_, v_n_) to v_0;kMer_ and leave the endpoint v_n_ undefined for the moment. We note that it is possible to have multiple kMer-edges carrying the same kMer as label (if the underlying subpaths are different). The edge probability distribution for the start node of the kMer-PRG is induced by the probabilities of the corresponding subpaths in the PRG. After creating edges for all subpaths emanating from v_0_, we compute the set of required vertices for the next level of the kMer-PRG, and attach the created edges to those vertices. We give the precise algorithm under point “C)”. This completes the kMerification of level 0 of the PRG.
2. To kMerify levels i = 1 … (L – k) of the PRG.

a. For each vertex v of l(v) = i in the PRG:

- For each kMer-edge e = (v_x_, v_y_) in the set K(v): Informally, the set of kMer-edges emanating from node v_y_ in the kMer-PRG is determined by how we can extend the subpath underlying e. Formally, the subpath underlying e has a defined last vertex v_z_ (v_z_ is a node of the PRG). We carry out a forward search for subpaths of length 1 starting at v_z_. Each subpath defines one possible extension for e and thus induces the creation of a kMer-edge e_x_ (as in step “A)”, the endpoint vertices for these edges remain undefined for the moment). The subpath underlying e_x_ consists of the subpath underlying e with its first symbol removed and extended by the subpath (of length 1) that induced the creation of e_x_. The label for e_x_ is generated by computing the kMer induced by the subpath underlying e_x_. (In particular, this means that the edge will be labeled with the “gap kMer” if the first symbol of the underlying subpath is a gap). The edge probability distribution at v_y_ is induced by the conditional probability to follow the found subpaths, conditional on being at node v_z_.
b. For the set of all kMer-edges created during the previous step “a”: Compute the set of required vertices for the next level I + 1 in the kMer-PRG, conditional on the set of kMer-edges created during the previous step, and attach the edges to their corresponding endpoint vertices (so far, these have remained undefined). We give the precise algorithm under point “3)”.
3. We describe how to compute the set of required vertices for a level i + 1 of the kMer-PRG, based on the set E_i_ of kMer-edges emanating from nodes of level i of the kMer-PRG. The result of this step (as explained above) will be that all edges that have “nearly” identical corresponding subpaths in the nucleotide PRG will end in the same node, with “nearly” here meaning that they are identical in all steps but the first. Each e = (v_m_, v_n_) ∈ E_i_ has an associated subpath in the PRG of the form (e_1_,e_2_,e_3_..). We define the m1-subpath of a subpath (e_1_,e_2_,e_3_..) as (e_2_, e_3_..), i.e. equal to the original subpath without the first edge. Clearly, there is a set of m1-subpaths induced by the set of subpaths associated with the kMer-edges in E_i_. We will create one vertex v at level l + 1 of the kMer-PRG for each unique m1-subpath (e_2_, e_3_..), and attach each kMer-edge to the vertex corresponding to the m1-subpath of its subpath. We note that this means that the original subpaths of the edges that end up attached to one vertex are identical from the second position onwards. We will thus attach <u>one</u> of the edges attached to the same vertex to K(v_x_), with the m1-subpath of the edge being of the form (e_2_, e_3_..) and e_2_ = (v_x_, v_y_).
4. As a post-processing step, mark all kMer-edges for which the underlying subpath includes edges that were marked as parts of the canonical reference sequence (see post-processing for the PRG construction algorithm). We also remove all subpaths that both consist exclusively of edges labeled with the wildcard character (“*”) and that can be removed without affecting the reachability of any subpath that is not exclusively labeled with wildcard symbols.

### 2.2 Edge compression

Edges in the kMer-PRG are labeled with single kMers. There are regions in the graph where each vertex has exactly one outgoing and one incoming edge, i.e. regions which consist of sets of non-branching subpaths. In a final transformation, we “compress” all such regions of the kMer-PRG into single edges, which are then labeled with (unordered) sets of kMer counts. We call the resulting graph the multi-PRG (“multi” because edges can be labeled with multiple kMers).

Formally, the definition of a multi-PRG is equivalent to the definition of kMers-PRGs, with the exception that each edge is now labeled with sets of the form {(kMer_1_, count_1_), (kMer_2_, count_2_), …}, where kMer_1_, kMer_2_ .. are unique kMers and count_1_, count_2_ .. specify how often the associated kMer is emitted upon edge traversal (we call these edges “multi-edges”).

Constructing a multi-PRG from a kMer-PRG is trivial. We give a short description of the algorithm.

1. Transform the kMer-PRG into a structurally equivalent multi-PRG, simply by substituting all edge labels *kMer* with {(*kMer*, 1)}.
2. Determine which regions from level i_x_ to level i_y_ in the multi-PRG satisfy the following criteria:

a. All nodes at level i_x_ have exactly one outgoing edge
b. All nodes at level i_y_ have exactly one incoming edge
c. All nodes with level > i_x_ and < i_y_ have exactly one outgoing edge and one incoming edge. This implies that there is one and only one subpath connecting each node from level i_x_ with a node from level i_y_ (and vice versa).
3. For all such regions from level i_x_ to level i_y_:

a. For all vertices v_x_ from level i_x_:

- Determine the subpath connecting v_x_ to a node v_y_ from level i_y_
- Create a new multi-edge by forming the union (with counts) of the labels of all edges traversed by the subpath. The new multi-edge connects the first and the last node traversed by the subpath.
- Delete all edges traversed by the subpath and all vertices traversed by the subpath except for the first and the last vertex.
4. The level function becomes temporarily ill-defined during the previous steps. There are different ways to deal with this in implementations. We use a temporary relaxation of the definition of the level function (specifically, dropping the requirement that vertices connected by one edge need to be assigned to subsequent levels) and a post-processing step to create a definition-consistent level structure (informally, by re-counting levels from the beginning of the graph).
5. As a post-processing step, mark all multi-edges created from edges that were marked for being part of the canonical reference sequence.

### 2.3 HMM

Multi-PRGs can be transformed into Hidden Markov Models (HMMs). One key challenge is that kMers which appear as edge labels on multiple levels of the graph violate the assumption that emissions from different states of the HMM to be created are independent. We deal with this by removing those kMers.

We and others have previously described how haplotype graphs can be transformed into haploid and diploid HMMs (see [1] and [2]), and we will only give a short informal description of this process here: each edge of the haplotype graph becomes a state in the HMM, and the emission probability distribution for each state is based on the underlying edge’s label. The resulting HMM will have a level structure analogous to that of the graph. State transition probabilities for the HMM are induced by the probability to move from one edge (v_x_, v_y_) to another edge (v_y_, v_z_) in the graph, i.e. by the edge probability distribution at v_y_.

We note that a modified version of that transformation can be applied to multi-PRGs. The structural transformations (i.e. from edges in the multi-PRG to states in the HMM and the state transition probabilities) are identical to the original transformation, and we refer the reader to [1] for details.

We now define the emission probability structure of the HMM. Specifically, we define the emission probabilities of an HMM state s with underlying multi-edge e = (v_x_, v_y_). We assume that there is a coverage parameter α and an error rate parameter β. α specifies the expected haploid coverage on kMers that are present in a sample genome being sequenced, and β specifies the expected haploid coverage on kMers present in the multi-PRG which are not present in the sample genome being sequenced (We define how to estimate these parameters below).

1. Identify and remove from all edge labels kMers which

- occur in the label sets of edges emanating from more than one level of the multi-PRG
- occur in a region of a canonical reference genome not covered by the PRG
- fail plausibility-based checks (see below)
2. Compute the set O_i_ of all remaining kMers used in the edge label sets of edges emanating from any node at level l(v_x_) = i, excluding

- “gap kMer” kMers (as “gap kMers” represent the empty string, they cannot generate an emission)
- kMers with ambiguous symbols (this is a simplification we make on practical grounds and one of the reasons for why we try to eliminate those as far as possible). Each kMer *kMer* ∈ O_i_ has an observed sample count o(*kMer*). o(*kMer*) is calculated by counting the number of occurrences of the word *kMer* in the sequencing reads from a sample.
3. We define the observed sample counts o(*kMer*) ∀*kMer* ∈ O_i_ as the data we will model for all states at the level of state s.
4. The state s that we want to define an emission probability distribution for has a label, and that label is a set of the form {(kMer_1_, count_1_), (kMer_2_, count_2_), ..}. We note that each kMer in O_i_ either appears in the label set for e, or not. If it appears in the label set for O_i_ as (*kMer*, *count*), we model the observed count o(*kMer*) with a Poisson distribution, with expected value α x *count* . If it does not appear in the label set for O_i_ (i.e. it is an error if we assume that s is the underlying state and the multi-PRG is a faithful representation of genomic variation), we model o(*kMer*) with a Poisson distribution with expected value β.

#### 2.3.1 kMer count plausibility checks

In the ideal case, the PRG is a comprehensive representation of genomic variation. However, in practice it is likely that there is genetic variation which is not captured by the PRG, which may distort our statistical approaches.

One important class of potentially confounding uncaptured variation is sequence duplication. In making inference, we generally exclude all kMers which occur in other parts of the reference genome or in other levels of the multi-PRG. Sequence duplication can have the effect that a kMer which we believe to be exclusively originating from one level of the multi-PRG is also present in another genomic location. That is, observed coverage on such a kMer not only comes from the multi-PRG, but also from other uncharacterized sources. One important symptom of this happening is excessive observed coverage on a kMer. Hence, for each kMer, we

- assume that observed coverage in read data is modeled by a Poisson distribution, with mean proportional to general kMer coverage and the kMer’s underlying genomic count
- assume a uniform prior on the kMer’s underlying genome count (ranging from 0 to at least 2 x the maximum number of occurrences of the kMer in any state of the multi-PRG)
- combine prior and Poisson model to obtain a posterior probability of underlying genome count for each kMer
- sum over all values of that posterior between 0 and ([maximum count of that kMer in any state of the multi-PRG] +2)
- exclude the kMer if that sum is < 0.5.

#### 2.3.2 Simplification

It is not uncommon to observe levels with thousands of states in the haploid HMM, and the number of states at a particular level increases by power 2 in the diploid version of the HMM (see [1] for the algorithm for creating a diploid version of the HMM).

We employ a simplification algorithm prior to applying the model to sample data: before creating the diploid HMM, we examine all edges of the multi-PRG. For each edge, we determine the fraction of kMers specified (weighted by counts) with no coverage in the sample (i.e. o(*kMer*) = 0). If this proportion is ≥ 0.5, we delete the edge – unless it was marked for being part of the canonical reference genome. (The rationale for this is to always have at least one remaining path even if coverage is 0. The canonical reference genome is thus the fallback path for the model.)

#### 2.3.3 Estimation of α and β

We estimate α, the coverage parameter, and β, the error rate parameter, from the multi-PRG.

For estimating α, we identify levels of the multi-PRG at which there is only one edge defined per level. If the multi-PRG faithfully reflects the underlying genome, all of these edges have to have diploid coverage. We sum over the observed sample coverage of the kMers specified by these edges and divide them by the sum of the edge-label specified kMer counts and a factor of 2.

For estimating β, we employ a simple greedy heuristic algorithm. We select (typically ∼ 200) levels of the multi-PRG according to these criteria:

- There are exactly two defined edges
- Both edges have the same number of non-gap non-ambiguous kMers (the label set element (*kMer*, *count*) contributes *count* to the kMer count of an edge).

We take α as estimated previously and start with an initial value for β (typically a small value below 1%).

At each selected level individually, we fit a simple Poisson model allowing for the three diploid “genotypes” defined by the two haploid edges (we use the same model as for the HMM emission probabilities, described earlier; this is conditional on current values of α and β). We select the maximum likelihood (ML) “genotype” call and treat coverage on edges not covered by the ML call as error.

After application to all selected levels, we sum over the total coverage on kMers not present on the edges of the ML calls, and divide it by the total number of kMers not present on these edges (i.e. a rate specifying how much coverage the kMers on edges which are likely not in the underlying genome receive). This gives us a new estimate for β. We repeat the estimation procedure, now conditional on the updated value of β, until the improvement in “total likelihood” of the data (here defined as the product of the likelihoods of the per-level ML calls) from one iteration to the next falls below a threshold.

### 2.4 Chromotypes

#### 2.4.1 Path inference

Standard statistical algorithms for HMMs can be applied to the defined HMM. Specifically, the Viterbi algorithm enables inference of one (diploid) Maximum Likelihood path through the model, and the Forward algorithm can be used to sample from the posterior distribution of paths [3].

The inferred diploid path is initially based on the multi-PRG, but can be mapped back unambiguously to equivalent traversals of the kMer- and nucleotide PRGs (each state of the [diploid] HMM is equivalent to two edges [one for each haploid traversal of the PRG] in the multi-PRG, and the transformation process from nucleotide PRG to multi-PRG can be reversed).

By concatenating the symbols emitted by the edges traversed by the ML diploid path through the nucleotide PRG, we create two “personalized reference haplotypes”.

However, the inferred diploid ML path (and hence the personalized reference haplotypes) loses phase at positions at which the underlying haploid HMM states are identical (that is, at positions at which both haplotypes traverse the same multi-PRG edge). “Loses phase” means that another diploid path, generated from the original path by flipping the two contained haploid paths after a position at which phase is lost, is statistically identical to the original path. For each Viterbi diploid path, we have thus a set of equally good diploid traversals which are identical to the Viterbi path in the edges that they traverse at each level, but not in haplotypic phase between different levels.

#### 2.4.2 Chromotypes

Chromotypes are a data structure to represent chromosomal genotypes at different levels of haplotypic resolution. The formal definition of a chromotype is that it is a PRG with not more than two edges at each level (as shown in the pictures below).

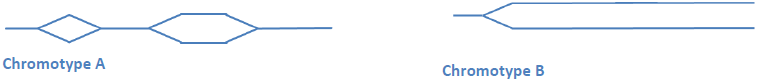

A diploid genome can be represented as two independent, completely resolved paths (“Chromotype B”), or as a sequence of homozygous stretches and heterozygous bubbles (“Chromotype A”) with phase lost whenever the chromotype enters a bubble – or as a mixture of the two approaches.

Chromotypes are a data structure well-suited to represent the set of Viterbi-equivalent diploid PRG traversals:

- Based on the state sequence of the diploid HMM, compute the equivalent two paths through the multi-PRG, and then the equivalent two paths through the kMer-PRG.
- Each path (traversal) if a sequence of nodes and edges through the kMer-PRG.
- Create a (kMer) chromotype by combining the nodes and edges (and their connectivity) from the two traversals into one new graph. Nodes and edges appearing in both paths will appear only once in the chromotype. If an edge or node appears in both haploid paths, it is to appear only once.
- If desired, this (kMer) chromotype can be converted to a nucleotide chromotype, for every kMer edge is equivalent to a subpath through the nucleotide PRG.

Chromotypes so-generated are subgraphs of the original PRG, describing an individual’s chromosomal genotype.

Chromotypes can also be created from sets of aligned strings. In the pipeline described here, this happens after read re-mapping to the Viterbi chromotypes – we use the re-mapped reads to identify new variants not yet represented in the PRG, but the discovery pipeline cannot deal with diploid reference genomes. Hence, we map to the personalized reference haplotypes, and modify the two strings according to the variants discovered. As a final step, we recombine the two modified haplotype strings back into a chromotype.

To create a (nucleotide) chromotype from two aligned strings, begin with a chromotype that has one start vertex and two completely resolved branches (encoding the two aligned strings) without branches. At each level, fuse nodes and edges of the two branches if the next k edge labels are identical for both branches.

#### 2.4.3 VCF creation

We note that the two personalized haplotypes imply a genotype at each position of the original MSA. It is sometimes desirable to express this information in VCF format. We give a sketch of the heuristic that we use to express one personalized reference haplotype in VCF format (the diploid case with two haplotypes follows then immediately).

There is a canonical reference which the VCF will refer to, also present as a row in the MSA used to create the PRG (in our case, this would be the chromosome 6 xMHC reference sequence “PGF”). At each column in the MSA, the canonical reference will either carry a specified non-gap character or a gap symbol.

For positions in the MSA at which the canonical reference is non-gap, creating the VCF for an inferred genotype is trivial. This includes cases in which the genotype to be expressed is a gap, but the canonical reference is non-gap – this means that the genotype to be expressed carries a deletion with respect to the reference, and all that is necessary is to find the starting position of the deletion with respect to the canonical reference. This is usually the first non-gap position to the left of the deletion in the haplotype to be expressed in a VCF, at which the canonical reference is also non-gap.

Positions where the haplotype to be expressed is non-gap, but the reference is gap, represent insertions in the haplotype with respect to the reference. In VCF format, insertions are usually expressed as longer alternative alleles at the position before the first inserted nucleotide. We can thus find the correct position for specifying an insertion by finding the first position to the left of the inserted sequence at which the reference row of the MSA is non-gap.

When creating VCFs, our algorithm produces additional files and annotations, which specify the positions in the MSA a variant in the VCF refers to.

## 3 Novel variant detection

### 3.1 Mapping reads to the personalized reference

After inferring two “personalized reference” haplotypes, using sequence read data from an individual, it is often desirable to use classical read-mapping tools to map the reads to the two personalized haplotypes. This enables the discovery of additional variants not yet present in the PRG, and can also be used as an additional step of quality control (regions of the personalized haplotypes with no coverage are generally less likely to faithfully represent the underlying genome than regions with good coverage).

We apply the following pipeline steps:

1. Create two “personalized reference genomes” by excising the region covered by the PRG and inserting the inferred personalized reference haplotypes.
2. Map read data from an individual independently to both personalized reference genomes. Mate-pair information could be used in this step. Our current implementation, however, ignores mate-pair information, for it cannot be assumed that the two inferred reference haplotypes correctly reflect long-range phase.
3. For each read mapped to the region covered by the graph, make a decision as to what personalized reference haplotypes it is likely to correspond to. We employ the following heuristic:

a. If a read is mapped to the region covered by the graph in only one personalized reference genome, assume that the read comes from the corresponding haplotype. As the rest of the reference genome remains unchanged, this implies that the mapping quality of the read over the region covered by the graph is higher than that of the read over alternative locations.
b. If the read maps to the region in both personalized reference genomes, use the mapping quality produced by the alignment software (and possibly other criteria like edit distance between read and the employed reference genome) to decide which of the two haplotypes the read should map to.
c. In cases of equal fit, choose uniformly.
4. Create two BAM files, covering the two personalized reference sequences and containing the reads mapped to each haplotype (from step 3).
5. Apply a variant-calling algorithm to the two BAM files, resulting in two VCF files.
6. Modify the two personalized reference haplotypes according to the produced VCF files (see below).
7. Finally, produce a VCF file representing the two modified personalized reference haplotypes (see Section “VCF creation”) and the corresponding chromotypes (see Section “Chromotypes”).

### 3.2 VCF-based reference haplotype modification

Step 6 of the algorithm specified above results in two VCF files, and we want to use the information from these VCF files to modify our personalized reference haplotypes.

We have developed a heuristic that solves this task to satisfactory degrees of accuracy (see the validation results presented in the main paper). However, we recognize the limitations of our approach. On a fundamental level, existing read mapping and variant calling tools cannot deal with diploid references, and this leads to complications in downstream analyses of the results.

The VCFs we want to use to modify the personalized haplotypes may specify insertions, deletions and single-nucleotide differences. We note that the two personalized reference haplotypes live in the coordinate space defined by the MSA. For each position in the MSA, we will now compute a set of implied novel genotypes (that is, implied by the VCFs based on the personalized reference haplotypes). We note that we need to allow for multi-character alleles in the columns of the MSA – to deal with insertions implied by the VCFs, for which there might be “no space” in the existing MSA coordinate system. The algorithm described under “VCF creation” can deal with multi-character alleles with small modifications.

We now describe how to integrate variants of different types into the two haplotypes. For each variant, the VCF it comes from will specify a position that it begins at; this position is relative to the personalized reference used a base for mapping, and can thus be translated into MSA coordinates. Note that each VCF will specify two alleles for each position; we will thus end up with up to four specified alleles at each position.

- Single-nucleotide differences: add the implied alleles to the set of implied alleles in the corresponding column of the MSA.
- Deletions: add the implied alleles (i.e., gap characters) to the set of implied alleles in the corresponding column of the MSA.
- Insertions: it might be that our original personalized haplotype called a deletion where there really is no deletion; and the VCF we analyze might reflect this by specifying an insertion at that position. We want to make sure that we place the characters specified by the insertion in the right columns of the MSA. Thus,

- Find the beginning of the insertion and the first character after the insertion, <u>relative to the personalized haplotype</u>, in the MSA.
- If there are no gaps between these characters, simply amend the column relating to the beginning of the insertion with the inserted characters.
- If there are gaps between these characters, temporarily fill the gap characters with the characters that the canonical reference specifies at these positions; use the Needleman-Wunsch algorithm [4] to align the canonical reference characters to the inserted allele; use this alignment to map each inserted character to a position in the MSA.

In the algorithm described, we have made no attempt to determine whether novel alleles should be integrated into the first or the second personalized reference sequence. We use a heuristic that tries to integrate variants in the haplotype that led to the generation of the variant-implying VCF; in most cases, however, the novel variants are SNPs and there is no (locally phase-determining) information to determine the haplotype of origin.

### 3.3 Current components

The above description of the algorithm does not assume the utilization of particular read mapping or variant calling tool. In our current implementation, we use BWA [5] for mapping and Platypus for variant calling. We also use samtools [6] for some of the intermediate steps.

## 4 Graph alignment and validation

### 4.1 A mapping algorithm for PRGs

The aim in conventional sequence alignment is to identify regions of homology (or more generally, similarity, according to a score function) between two sequences. In the context of the analysis of next-generation sequencing data analysis, sequence alignment (or mapping) between the sequence of a read and the reference genome is used to identify the putative genomic origin of the read.

The alignment problem exists for PRGs, too: identify regions of homology between a given sequence (henceforth called “query sequence”) and a population reference graph. There are two challenges. First, PRGs specify a set of possible paths through the graph, and the query sequence could align to any of these paths. Second, PRGs can comprise “gap” edges – so the optimal alignment would specify “gap” symbols whenever a “gap” edge is traversed, but without penalizing them in the way normal gaps in the alignment are typically penalized.

To address these challenges, we have developed a generalization of the Needleman-Wunsch algorithm for global sequence alignment. We have also developed a set of approximations to reduce the computational complexity of the algorithm, enabling the alignment of query sequences on the order of thousands of bases to PRGs on the order of millions of levels (i.e. regions or genomes Megabases in size).

#### 4.1.1 Statement of the problem

There is a query sequence Q=(q_1_,…,q_N_Q__) consisting of N_Q_ characters from the alphabet {“A”, “C”, “G”, “T”} and a PRG G = (V, E).

Define P_sub_ as the set of possible subpaths (v_a_,v_b_)(v_b_,v_c_)…(v_m_,v_n_) through the graph (for definition of “subpath”, see Section “kMerification”) and let P_traversal_ contain all elements of P_sub_ for which v_a_ is the start vertex and v_n_ is one of the final vertices. Informally, the set P_traversal_ contains all complete traversals of the graph.

Define a symbol g_2_ which is neither contained in the query alphabet nor exists in E or as a label of one of the edges in E. g_2_ denotes the “gap” symbols introduced during the alignment process, which will be penalized, and, importantly, it is by definition different from the “gap” symbols already contained in G.

We call A = (Q’, E’) an **alignment** of query sequence Q to PRG G of length A_L_if

1. A = (Q’, E’) = ((Q’_1_, Q’_2_, ..,Q'_*A*_*L*__), (E’_1_, E’_2_, .., E'_*A*_*L*__)), with
2. Q’_x_ ∈ {“A”, “C”, “G”, “T”, g_2_} ∀x ∈{1 … A_L_}, and
3. E’_x_ ∈ E ∪{g_2_} ∀x ∈{1 … A_L_}
4. Q’ can be formed by inserting an arbitrary number (including 0) of g_2_ elements into Q
5. E’ can be formed by inserting an arbitrary number of g_2_ elements into an element of P_traversal_
6. There is no position x in the alignment with Q’_x_ = E’_x_ = g_2_.

For a given PRG and a given query sequence Q, define A_all_(Q, G) as the (finite) set of possible alignments,

The **alignment problem** is to maximize a score function SCORE(*A*) → ℝ on the set of possible alignments A_all_(Q, G).

#### 4.1.2 Scoring

Alignment scoring functions SCORE(*A*) → ℝ for A ∈ A_all_(Q, G) can assume arbitrary form (we require that higher scores for a given pair A, G indicate better alignment quality, i.e. we want to maximize SCORE to find a good alignment).

From standard sequence alignment, however, it is well-known [7] that scoring functions of certain forms are amenable to efficient optimization via dynamic programming, in particular those which assign a fixed score to every column in the alignment with identical characters (“matches”), a fixed score to every column in the alignment with non-identical, non-gap characters (“mismatches”), and either a fixed or a linear score to every column containing gap characters. In the linear gap score case (“affine gap penalties”), the first gap in a sequence typically gets a lower score (“gap opening penalty”) than all subsequent gaps in the same sequence (“gap extension penalty”).

We will now define equivalent scoring functions for graph alignment. We define the function label(x) as the label of edge x if x is an edge, and as the empty string “” if x is equal to g_2_. We need to make special provisions for the case where label(e) for an edge e is the “gap” symbol from the PRG.

##### 4.1.2.1 Non-affine gap penalties

Let a score function SCORE(A) = SCORE(Q’, E’) assume the following form

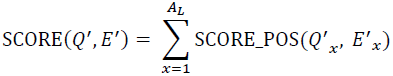

with

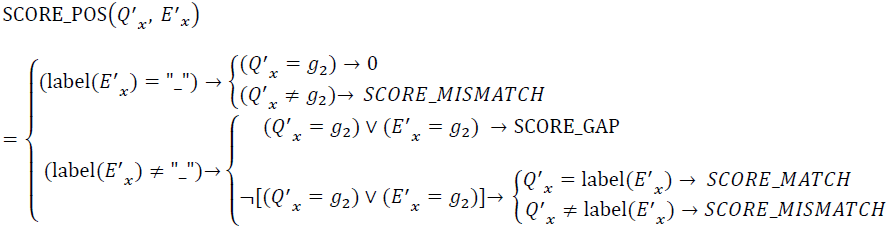

We note that we assign score 0 to columns in which the traversed edge carries the “gap” symbol and in which the corresponding sequence column carries the (alignment-induced) gap-symbol g_2_. Thus, the scoring function will neither reward nor penalize such columns, if SCORE_MATCH > 0 and SCORE_MISMATCH and SCORE_GAP< 0.

##### 4.1.2.2 Ends-free alignment scores

Ends-free alignment as defined here is typically applied if the query sequence is much shorter than the graph; the effects are (1) that the gaps which are necessary to extend Q to at least the graph’s length do not get penalized and (2) to favour “dense” alignments in which there are not many gaps between the original characters of Q.

Formally, in ends-free alignment, SCORE_GAP = 0 for position x if *Q*′*_x_* = *g*_2_ and ∀*x*_2_ ∈ {1 .. *x*}: *Q*′*_x_*_2_ = *g*_2_ or if *Q*′*_x_* = *g*_2_ and ∀_*x*_2__ ∈ {*x*.. *A*_*L*_}: *Q*′_*x*_2__ = *g*_2_.

##### 4.1.2.3 Affine gap penalties

In affine-gap alignment, the first gap in a continuous sequence of gaps is typically scored differently from the subsequent gaps. “Opening” a gap is typically associated with a higher penalty than “continuing” a gap. Traversing a sequence of “graph gaps” should not end an affine gap in query sequence space.

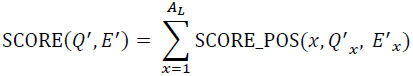

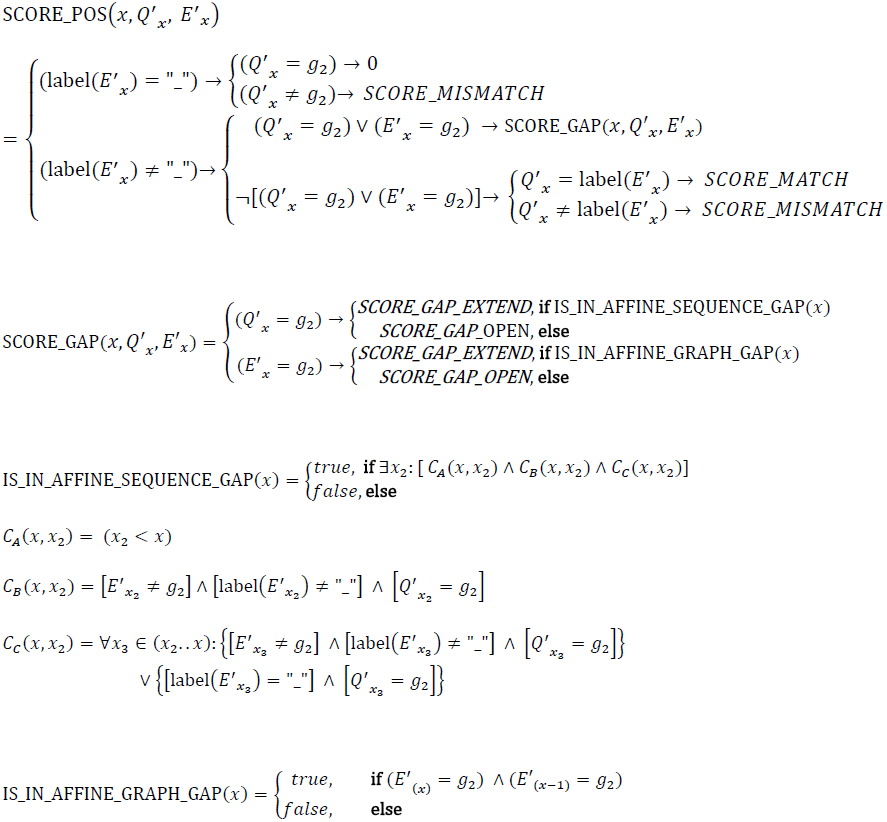

For x = 0, IS_IN_AFFINE_SEQUENCE_GAP(x) and IS_IN_AFFINE_GRAPH_GAP(x) are defined as *false*.

We summarize the effect of these scoring functions:

- For columns with two non-gap characters, we add either SCORE_MATCH or SCORE_MISMATCH.
- g_2_ characters in E’ are always penalized as gaps, and the first such character in a sequence typically more strongly so than the following characters.
- g_2_ characters in Q’ are only penalized as gaps if the corresponding character in E’ is not the “gap” character “_”. In this case (i.e. if the g_2_ character is penalized), it is determined whether this g_2_ character is the first one of an affine sequence gap, and a penalty is assigned accordingly. Affine sequence gaps can span columns in which E’ contains “_” symbols. In terms of the definitions made above, an alignment position *x* is only part of an affine sequence gap if conditions *C_A_*(*x*, *x*_2_), *C_B_*(*x*, *x*_2_) and *C_C_*(*x*, *x*_2_) are all true for *x* and a *x*_2_ < x:

- C_*A*_(*x*, *x*_2_): true if *x*_2_ < x
- C_*B*_(*x*, *x*_2_): true if at *x*_2_ a sequence gap is initiated, i.e. there is a gap in the query sequence but no gap in the graph sequence at that position
- *C_c_*(*x*, *x*_2_): true if the gap initiated at *x*_2_can be extended to *x*, i.e. all graph alignment positions in between carry either a defined graph character or a “_” graph gap symbol, and all sequence alignment positions in between carry a gap symbol.

Note that cases in which both Q’ and E’ carry the g_2_ character at a column are also invalid by definition.

#### 4.1.3 Maximization

The solution to the alignment problem is found by maximizing the supplied scoring function over the set of all possible alignments A_all_(Q, G).

For the scoring functions described above, the maximization can be carried out by a dynamic programming algorithm, very similar to the Needleman-Wunsch algorithm.

The main difference is that the Needleman-Wunsch algorithm utilizes a two-dimensional matrix of scalars, whereas we utilize a two-dimensional matrix of vectors. For the Needleman-Wunsch algorithm, the value in cell (x,y) of the scoring matrix is defined as the maximum score attainable after having consumed x characters from the reference sequence and y characters from the query sequence. For our algorithm, each cell carries a vector, and we index the values in this vector by a third coordinate. Value (x, y, z) is then defined as the maximum score attainable after having consumed x levels from the graph, y characters from the query sequence, and ending up in vertex z of level x of the graph.

##### 4.1.3.1 Computation for non-affine gap penalties

PRG G has L levels, and query sequence Q has N_Q_ characters.

At each level l of G, let Z_l_ be the number of vertices at this level. Use the integers 1 .. Z_l_ to arbitrarily enumerate the vertices of each level l, and define two index functions that map between vertices and their associated indices and vice versa:

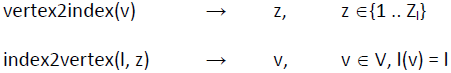

Now define a (N_Q_ + 1) × L matrix denoted M. Each cell (q_i_, l) contains a vector of length Z_l_. (We use l to index levels of the graph, and we use q_i_ to index positions within query sequence Q). We use the notation M(q_i_, l, z) to denote the z-th value of the vector in cell (q_i_, l).

For the purpose of this section, we use **0-based** indices to index M, the graph and Z_l_. For example, Z_0_ refers to the number of vertices at the first level of the graph. Individual vertices, however, are indexed using 1-based indices, i.e. the first vertex at a level has the index 1.

We define auxiliary set functions that map a vertex to its potential ancestors and potential predecessors, along the graph.

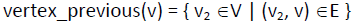

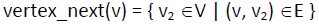

By definition, the previous and next vertices for v are one level below / one level above v.

We will fill M cell-by-cell. (q_i_, l, z) shall be the maximum attainable score after having consumed q_i_ characters from Q, l levels from G and ending up in the z-th node at level l of G.

We initialize M:

- Origin: For all z∈ (1 .. Z_0_), set M(0, 0, z) = 0.
- Gaps along the graph coordinate: For all l ∈ (1 … [L – 1]) and z ∈ (1 .. Z_l_), set

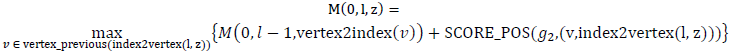
- Gaps along the query sequence coordinate: For all q_i_ ∈ (1 … N_Q_) and z ∈ (1 .. Z_0_), set

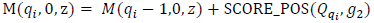

We progressively fill M using a nested loop, in the order specified:

For each l ∈ (1 … [L – 1]), for each q_i_ ∈ (1 … N_Q_), for each z ∈ (1 .. Z_l_):
Define *v*_*P*_ = vertex_previous(index2vertex(*l*, *z*)).

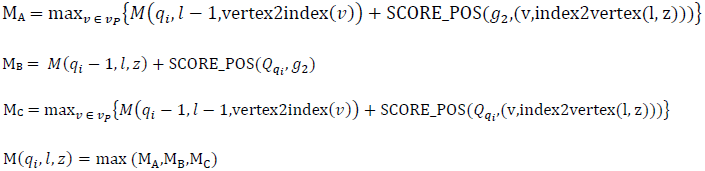

(If there are pairs of vertices connected by more than one edge, the maximization in M_A_ and M_C_ needs to be carried out explicitly over edges, instead of nodes at the previous level).

In this recursion, M_A_ is the “gap in query sequence” step, M_B_ is the “gap in graph” step, and M_C_ is the “match or mismatch” step.

The final maximum is max_z ∈{1..*Z*_*L*−1}__ *M*(*N*_*Q*_, *L* − 1, *z*), and backtracking, analogous to the classical Needleman-Wunsch, will identify the corresponding alignment. This is easily implemented by having a second matrix, in dimensionality equivalent to M, which stores which coordinates each maximum was drawn from.

##### 4.1.3.2 Computation for affine gap penalties

In classical sequence alignment, affine gap penalties are realized by progressively filling three matrices: one matrix for paths that end in an affine “query sequence” gap, one matrix for paths that end in an affine “reference sequence” gap, and finally one matrix for paths that end arbitrarily, including those that end with a match or mismatch. Jumps between these matrices are allowed where appropriate according to these definitions. For example, the third matrix would always contain the maximum value of all three matrices for any given coordinate. We shall now proceed accordingly, but extend the classical framework to deal with graphs.

We use all definitions from the previous section where appropriate but instead of M, we now define three matrices M_D_, M_G_ and M_S_, all of dimensionality (N_Q_ + 1) × L. As for M, each cell in these matrices contains a vector with as many elements as there are nodes at the corresponding level of the graph.

M_G_ (q_i_, l, z) shall contain the maximum attainable score of all paths ending in an affine graph gap (i.e. a gap *in between levels of the graph*, consuming a character of the query sequence), after having consumed q_i_ characters from Q, l levels from G and ending up in the z-th node at level l of G. M_S_ (q_i_, l, z) shall contain the equivalent score for paths ending in an affine sequence gap. Finally, M_D_ (q_i_, l, z) shall contain the equivalent score for paths ending arbitrarily.

We note that we need to make sure that the algorithm implements the provisions we specified in Section “Scoring” for edges carrying the graph “gap” symbol. Importantly, affine sequence gaps can go “through” such edges, but they must not begin by traversing them.

Initialization:

- Origin: For all z∈ (1 .. Z_0_), set

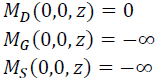
- Sequence gaps (along the graph coordinate): For all l ∈ (1 … [L – 1]) and z ∈ (1 .. Z_l_): Define:

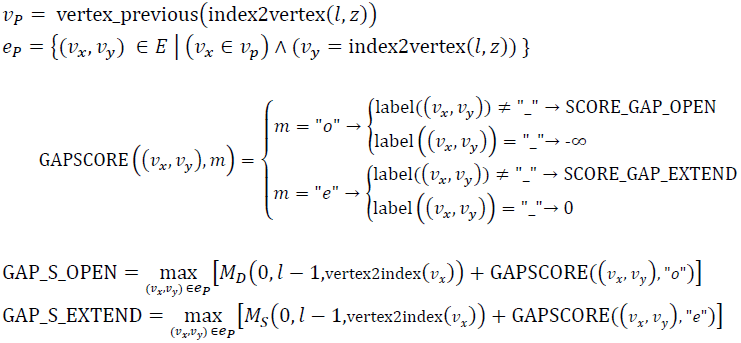 Set:

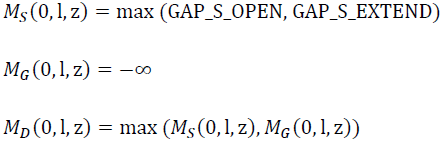
- Graph gaps: For all q_i_ ∈ (1 … N_Q_) and z ∈ (1 .. Z_0_): Define:

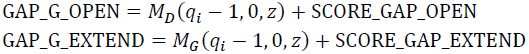 Set:

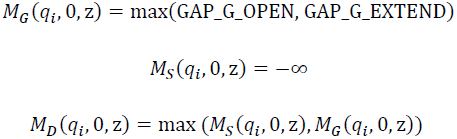

We progressively fill the three matrices using a nested loop, in the order specified: For each l ∈ (1 … [L – 1]), for each q_i_ ∈ (1 … N_Q_), for each z ∈ (1 .. Z_l_):

**M_S_:**
Define:

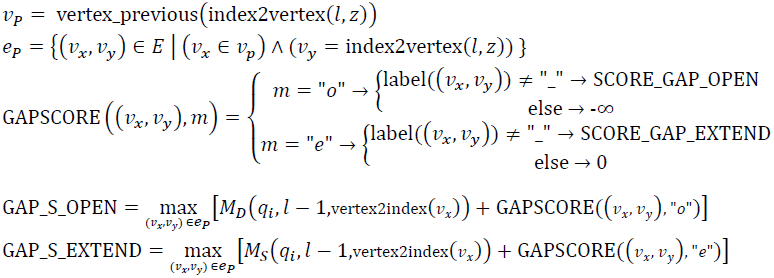
Set:

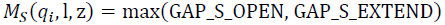
**M_G_:**
Define:

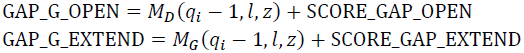
Set:

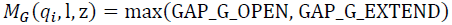
**M_D_:**
Define:

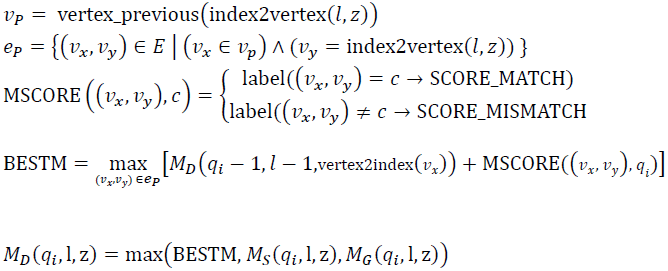

The final maximum is max_*z* ∈{1..*Z*_*L*-1}__ *M*_*D*_(*N*_*Q*_, *L* – 1, *z*), and backtracking will identify the corresponding alignment. For the affine-penalty algorithm, it is necessary to keep track not only of coordinates but also of the movements between the matrices.

#### 4.1.4 Parameterization

In our implementation, we use the following parameterization:

SCORE_MATCH = 2

SCORE_MISMATCH = -5

SCORE_GAP_OPEN = -4

SCORE_GAP_EXTEND = -2

#### 4.1.5 Implementation

The complexity of the described algorithm in in the class O((N_Q_ + 1) × L x max(Z_l_ × Z_(l-1)_)) – i.e. practically inapplicable to problems of the scale we are most interested in: PRGs with millions of levels and query sequences ranging from hundreds to tens of thousands of nucleotides.

We have thus developed a “seed and extend” approximation to the full algorithm, the key components of which we outline here.

Informally, we utilize stretches of sequence we can uniquely localize to constrain the alignment search space. Each such uniquely localized stretch relates to a particular subpath through the full alignment matrix, i.e. a defined combination of matches and gaps connecting one particular point in the alignment matrix with a second particular point in the alignment matrix. In order to complete the alignment, we need to connect these subpaths (a) to each other and (b) to the top-left and bottom-right corners of the matrix, at which point all graph levels and query sequence characters will have been incorporated.

##### 4.1.5.1 Step 1: Chaining

We scan through the query sequence from left to right and identify all exact and contiguous matches between subpaths in the PRG and the query sequence. Each subpath we refer to as a “chain”.

To speed up this process, we kMerify the PRG that we map to, keeping track of the subpath spanned by each kMer. We store kMers and corresponding subpaths in a hash table.

We require that each chain begin with an exact kMer match, and we extend each chain until we hit a mismatch. Of note, each kMer can initiate multiple chains.

##### 4.1.5.2 Step 2: Global chain filtering and fixing

Each chain specifies a path through the main scoring matrices. If the divergence between query sequence and PRG is not too high (which we assume as the PRG / chromotypes we map to contain many population / individual variants), connecting and extending these chains in a sensible manner should yield a good alignment.

We rate each chain by kMer double-uniqueness. That is, for each kMer we determine whether it occurs exactly once in the query sequence and exactly once in the PRG we map to. If both conditions are satisfied, we say that the kMer is double-unique. For each chain we determine the proportion of double-unique kMers, and we rank the chains according to this criterion. We store the ranked chains together with their proportion of uniqueness in a list structure we call `AVAILABLE_CHAINS`.

If the mapping algorithm is run in deterministic mode, we carry out the following steps:

1. Determine whether there are still chains in `AVAILABLE_CHAINS` (optionally meeting the criterion that the absolute number of double-unique kMers per chain is above a certain threshold – we currently use 1) – if not, terminate.
2. Select the highest-ranked chain from `AVAILABLE_CHAINS` and store the corresponding alignment matrix subpath (i.e. the induced sequence of matches and gaps).
3. Remove the selected chain from `AVAILABLE_CHAINS` and remove all other chains from that set which are incompatible with the selection. Incompatibility can be induced 1) by basic alignment structure as well as 2) by the properties of the PRG we align to. For the first point, consider fixing a chain which maps the first 10 kMers of the query sequence to levels 10–15 of the graph. It is clear that no following kMer (to the right of the ones already fixed) can be mapped to levels 1–9 of the graph. For the second point, consider that not all edges in the PRG are necessarily reachable from all other edges, even if they are compatible in terms of levels. PRGs can, for example, contain long haplotypic paths with no connecting edges. If a fixed chain maps to the first of two such paths, no other chain can map to the second.
4. Go to step 1.

If the algorithm is run in probabilistic mode, step 2 is replaced with a probabilistic selection, the chains weighted by optimality.

##### 4.1.5.3 Step 3: Recursive local chain filtering and fixing

We now deal with the “squares” in between the fixed chains from Step 2 (and the area between the origin of the alignment matrix and the first chain, and between the last chain and the bottom-right corner of the alignment matrix).

Each fixed chain from Step 2 has start- and endpoints, defined in terms of their (q_i_, l, z) coordinates. Each region between two chains from Step 2 has thus defined start- and endpoints with (q_i_, l) coordinates assigned, and we refer to these regions as “squares” (the z coordinates matter, too, but we nevertheless we stick with the two-dimensional metaphor).

We now apply the algorithm from Step 2 to each square so-defined, modifying the measure of kMer uniqueness to only take into account uniqueness within the square (along the q_i_ and l coordinates, i.e. we reduce our notion of uniqueness both in terms of levels of the graph and in terms of the query sequence).

This will typically enable us to fix more chains, and we recursively repeat this procedure for each square in between the new sets of chains until we can't fix any more chains.

##### 4.1.5.4 Step 4: Chain extension

We now use the global graph alignment (described earlier in this section) algorithm to try to fill the space in the remaining squares (squares between chains from Step 2 and Step 3).

More formally, for each remaining square, we start running the global alignment algorithm at the start coordinates and (in reverse direction) at the end coordinates. We use affine gap penalties with the parameterization described earlier. We terminate the algorithm if either the square boundaries have been reached or if the total score has fallen below a certain threshold (-11). The algorithm also differs from the one described earlier in a couple of other points, most of them aimed at eliminating unpromising areas of the search space:

- The scoring matrices are filled in diagonals. Each diagonal counts as one iteration, and the termination threshold refers to the maximum value achieved during the computation of one diagonal.
- We also measure how many iterations (i.e., filled diagonals) ago the achieved maximum value over the complete search space was last increased, and if this number crosses a threshold (40), we terminate.
- We measure the maximum value achieved in one diagonal, and we prune all cells in the diagonal if the difference between the maximum and the cell value (in M_D_) is bigger than a threshold (15). Pruned cells are not considered as sources for the recursion equations when computing the next diagonal.
- If the total maximum over the scanned area is achieved in multiple cells, we store all corresponding coordinates. If the reverse run started from the end coordinates hits one of the maximum points of the forward run started from the start coordinates, we also store the coordinates and the achieved score.

When the forward and backward extension runs have terminated, we have obtained

1. a maximum achieved score for each run and where this was achieved
2. (potentially) where the reverse run hit the maxima of the forward run, and the associated scores.

For notational convenience, let R_F_ denote the set of coordinates from (1) and (2) from the forward run, and let R_B_ denote the set of coordinates from (1) and (2) from the backward run. If the total maximum (1), however, is < 0 for a run, we will only include the start coordinates of the run in the corresponding set.

We now consider the cross product R_F_ × R_B_. For each combination of ends points {(*f*_*q*_*i*__, *f*_*l*_, *f*_*z*_), (*b*_*q*_*i*__, *b*_*l*_, *b*_*z*_)} ∈ (R_F_ × R_B_), we

- examine whether the combination of forward- and backward-derived coordinates is compatible, i.e. whether it is possible to connect the forward end point (*f*_*q*_*i*__, *f*_*l*_, *f*_*z*_) to the backward end point (*b*_*q*_*i*__, *b*_*l*_, *b*_*z*_) via a PRG-consistent alignment path in positive direction along the l and q_i_ coordinates. (This path is typically a long sequence gap followed by a graph gap, or vice versa, unless *f*_*q*_*i*__ = *b*_*q*_*i*__ or (*f*_*l*_, *f*_*z*_) = (*b*_*l*_, *b*_*z*_)). If there is no such path, we discard this combination.
- If there is such a path, however, we compute the score for

- the path from the forward extension start point (i.e. the top-left corner of the square) to the particular forward extension end point (*f*_*q*_*i*__, *f*_*l*_, *f_z_*)
- the path from (*f*_*q*_*i*__, *f*_*l*_, *f_z_*), to (*b*_*q*_*i*__, *b*_*l*_, *b_z_*)
- the path from (*b*_*q*_*i*__, *b*_*l*_, *b_z_*) to the backward extension start point (i.e. the bottom-right corner of the square).
- The three paths combined connect the start coordinate of the square to the end coordinate of the square, and the three scores combined determine the optimality of that particular way of connecting the coordinates. In summing up the scores, we need to keep track of where affine gaps open and close.

We finally select the combination that achieved the maximum score and use the corresponding path to connect the start coordinate of the square to the end coordinate of the square. (If multiple combinations achieve the same score, we make a random selection; if no combinations are compatible, we connect the start and the end coordinate of the square with gaps).

##### 4.1.5.5 Step 5: Backtracking

We choose max_z ∈{1..*Z*_*L*−1}__ *M*_*D*_(*N*_*Q*_, *L* – 1, *z*) as the final score of the algorithm, and we backtrack accordingly.

##### 4.1.5.6 Further points

- To speed up the algorithm, in particular for short or medium-sized query sequences, it can be helpful to omit the computation of the optimal gap paths between the top-left corner of the alignment matrix and the first chain and the last chain and the bottom-right corner of the alignment matrix. To see why this makes sense, consider aligning a fragment of 1000 bases against a PRG of 5m bases: In the resulting global alignment, approximately 4.999m positions would be used to specify the gaps before and after the query sequence, and computing these gaps would typically be much more resource-intensive than the parts relating to the query sequence. (Graph alignment requires traversal of the graph, even if a sequence gap is to be inserted at a particular position – for the final score is influenced by whether the sequence gaps sit below edges labeled with the “gap” symbol or not).
- The algorithm described in Step 2 (and hence that for Step 3) can, as specified, be run in probabilistic mode. We typically carry out one “deterministic” run (always selecting the chain with maximum double-uniqueness) and a number of “probabilistic” iterations (randomly selecting from a uniqueness-weighted selection of chains). As final result we select the iteration that achieved the highest score, and compare across iterations to compute measures of confidence. For example, for each character in the query sequence, we count in how many iterations it ends up being assigned to the same level of the PRG as in the chosen maximum iteration. This (to an extent) quantifies the uncertainty in placement of query characters.

### 4.2 Validation

#### 4.2.1 Chromotype disentanglement for kMer recovery validation

For any two (or more) positions at which a chromotype loses phase (i.e. at nodes with two outgoing edges) within distance k (where k is the chosen kMer length for validating the chromotype against some sequencing data), we need to disentangle the chromotype prior to validation.

After disentanglement, a chromotype induces a set of well-defined kMers, which we expect to find in sample sequencing data if the chromotype is a correct representation of sequence present in the sample.

We employ a simple greedy algorithm for disentanglement. As our final criterion for assessing a chromotype is how many of the kMers can be recovered from sample data, our disentanglement procedure (locally) optimizes for this criterion.

To prepare for disentanglement, we compartmentalize our chromotype so that all stretches between the phase breakpoints become one compartment (i.e. the compartments are separated by nodes with more than one outgoing edge). Each compartment can be either homozygous or heterozygous, depending on whether the stretch it spans has one or two nodes at each level.

Now we move through the chromotype from the left to the right, carrying with us a set of unresolved haplotype pairs. If the first compartment of the chromotype is homozygous with sequence s_1_, we initialize the set HAPSET of pairs of haplotypes as {(s_1_, s_1_)}, or as {(s_1_, s_2_)} if it is diploid with sequences s_1_ and s_2_.

Moving from compartment i to compartment i + 1, we carry out the following procedure:

- If compartment i + 1 is homozygous, append sequence s_1_of compartment i + 1 to all haplotype pairs in HAPSET. If s_1_contained k or more non-gap characters, call REDUCE. Set i = i + 1.
- If compartment i + 1 is heterozygous, append sequences s_1_ and s_2_ to all members of HAPSET, once matching s_1_ with the first member of each haplotype pair and once with the second. That is, HAPSET doubles in size. If HAPSET has more than 100,000 members, call RESOLVE. Set i = i + 1.

Before defining REDUCE and RESOLVE, we define OPTIMALITY((s_1_, s_2_)). OPTIMALITY computes the optimality of a haplotype pair (s_1_, s_2_) by

- Removing all gaps from (s_1_, s_2_).
- Dividing the number of kMers in (s_1_, s_2_) found in the sample data by the total number of kMers in (s_1_, s_2_).
- If there are 0 kMers in (s_1_, s_2_) (e.g. because both sequences have length smaller than k), OPTIMALITY is defined as 0.

RESOLVE orders HAPSET according to the values returned from OPTIMALITY and eliminates all members of HAPSET but the one with the best score.

REDUCE orders HAPSET according to the values returned from OPTIMALITY and retains the 1000 best-scoring haplotype pairs.

After completing this algorithm for the last compartment (we need to call RESOLVE if HAPSET has more than one member), the chromotype is disentangled into two strings (the members of the only remaining element of HAPSET), which specify an unambiguous set of kMers. We call the chromotype equivalent to these two strings (i.e. one start vertex and two separate, non-connected branches encoding the two strings) the disentangled chromotype.

#### 4.2.2 Identification of xMHC-specific contigs

For the Moleculo-based validation, we want to identify contigs that have originated from NA12878’s xMHC region. We thus filter and trim the raw contig sequence data prior to alignment, according to the following criteria:

- We compute the set of all kMers (k = 31) occurring in the kMerified xMHC PRG. We call all kMers occurring in this set “xMHC kMers”.
- We also compute the set of all kMers (k = 31) occurring in the human reference genome, excluding the region covered by the xMHC PRG. We call all kMers in this set “reference kMers”. Note that some kMers are both xMHC kMers and reference kMers. We call kMers which are xMHC kMers but not reference kMers “xMHC-unique kMers”.
- We filter contigs according to the following criteria:

- Fraction xMHC kMers >= 0.8
- There are two xMHC-unique kMers spanning a stretch of at least 50 bases (in between the two kMers). For each contig, we select the maximum stretch MAXSTRETCH spanned by two such xMHC-unique kMers.
- Within MAXSTRETCH, fraction of xMHC-unique kMers >= 0.5
- Within MAXSTRETCH, fraction of reference kMers <= 0.3
- If a contig passes these tests, we truncate the contig to MAXSTRETCH and align MAXSTRETCH.

### 4.3 Important symbols and abbreviations

A: Alignment A
Q=(q_1_,…,q_N_Q__): Query sequence of N_Q_ characters
A_all_(Q, G): Set of alignments between Q and G.
PRG: Population Reference Graph
COV: Catalogue of Variation

G: The specific PRG
V: Set of vertices
E: Set of edges
P_n_(e): Edge probability distribution at node n
P_sub_: Set of all subpaths
P_traversal_: Set of all subpaths, constrained to complete traversals
V_m_: Two vertices
v_n_: 
e: One edge
l(v): The level of vertex v

L: Scaffold haplotype MSA length; last level of haplotype graph
H(v): The set of scaffold haplotypes attached to v
K(v): The set of kMer-edges attached to v
c_v_: Current vertex

r: “Recombination” parameter
S_N_: Number of scaffold haplotypes
S_n,i_: i-th position (MSA) of haplotype n
O_i_: Set of kMers output from level i
o(kMer): Sample count of kMer *kMer*
x: Generic variable
X: Additional variant specifiers.

*suffix*(v, r): Suffix function for vertex v of length r

Q: Alignment query sequence
N_Q_: Length of Q
q_i_: Index for Q
Q’: Aligned query sequence
E’: Aligned edge sequence
A_L_: Alignment length
M: Alignment scoring matrix
Z_l_: Number of nodes at level
node(l, z): Retrieve node z at level l.

## 5 Experimental details

### 5.1 Simulation

We carry out simulations by independently generating two paths through the PRG (uniform choices at junctions) and treating these as a sample’s underlying diploid genome. We concatenate the edge labels induced by each path, remove “gap” characters and use the strings so-generated as a sample’s two haplotypes to generate reads from. At each position, the number of starting reads (read length 85 bp) is given by a Poisson distribution, parameterized to match an expected total coverage of 30x. Accuracy is assessed by comparing the diploid true underlying genotype at each level of the PRG with the diploid genotype induced by the Viterbi genotype computed from the simulated reads. Our simulations are limited in that a) we ignore read error (the main effect of which is a slight reduction of coverage) and b) we treat the simulated paths as a sample’s complete genome.

### 5.2 Graph for the extended MHC

We define the extended MHC (xMHC) as the genomic region spanned by the “PGF” xMHC haplotype (identical to the canonical human reference in the region – in B37 coordinates: chr6:28,702,185-33,451,429).

We use the eight xMHC haplotypes from the MHC haplotype project [8] as scaffold haplotypes for the region. We create an MSA for the eight haplotypes using the programs FSA [9] and MAFFT for refinement [10] . We use the SNPs identified by the 1000 Genomes Project, Phase 1, Release 3 [11] as additional variant specifiers for the eight MHC haplotypes.

We also use all available genomic HLA allele sequences from IMGT/HLA ([12], http://www.ebi.ac.uk/ipd/imgt/hla/, downloaded on 29/02/2012) for the classical HLA alleles at the loci *HLA-A*, *-B*, *-C*, *-DQA1*, *-DQB1*, *-DRB1* as additional scaffold haplotypes (these scaffold haplotypes cover all exons and introns of the genes – for many known alleles, the genetic sequences are not completely specified over all exons and introns, but the PRG construction algorithm we have defined removes most of the wildcard characters found at the unspecified positions). We do not specify any additional variant specifiers for the classical HLA genes.

The edge probability distributions we specify at each vertex in the PRG are mostly improper. Specifically, we assign probability 1 to each edge. This is motivated by the downstream parts of our pipeline: we mostly rely on the Viterbi algorithm for inferring Maximum Likelihood personalized haplotypes. With the specified improper parameterization, each possible path through the model is equally likely under the Viterbi algorithm, independent of how many potential branching points (vertices where there is more than one possible edge to follow) it contains.

We use kMer length k = 31 for creating the kMer-PRG.

#### 5.2.1 Ensembl inconsistency

In the process of examining available annotation information for the eight xMHC haplotypes, we discovered an inconsistency in the Ensembl database [13]. On the SSTO haplotype, *HLA-DRB1* and *HLA-DRB4* were mapped to the same start coordinate, likely caused, according to Ensembl, by a mismapping of exonic sequence of the two transcripts ENST00000549627 and ENST00000548105 (*HLA-DRB4* and *HLA-DRB1* exon sequence is similar). The two transcripts will be deleted in release 72 or release 73 (*personal communication*).

### 5.3 Sample details and HLA types

CS1 and CS2-6 samples:

Next-generation sequencing data generated as described in the main text. Sample read data available at request from GlaxoSmithKline.

NA12878:

Next-generation sequencing for NA12878 from the Illumina Platinum genomes project (www.illumina.com/platinumgenomes/) was downloaded from the EBI (www.ebi.ac.uk/ena/data/view/ERP001775). Sample data details described in the main text.

Sample HLA types (reported to 4-digit accuracy using ‘g’ nomenclature):

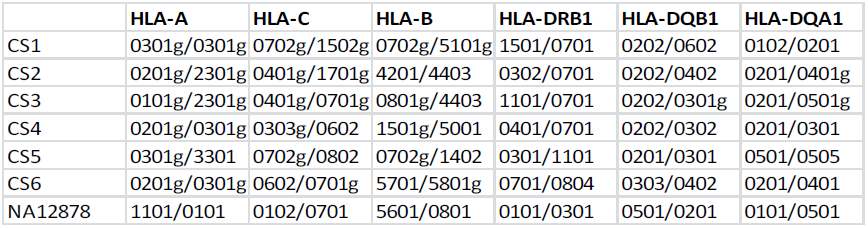

|  | HLA-A | HLA-C | HLA-B | HLA-DRB1 | HLA-DQB1 | HLA-DQA1 |
| --- | --- | --- | --- | --- | --- | --- |
| CS1 | 0301g/0301g | 0702g/1502g | 0702g/5101g | 1501/0701 | 0202/0602 | 0102/0201 |
| CS2 | 0201g/2301g | 0401g/1701g | 4201/4403 | 0302/0701 | 0202/0402 | 0201/0401g |
| CS3 | 0101g/2301g | 0401g/0701g | 0801g/4403 | 1101/0701 | 0202/0301g | 0201/0501g |
| CS4 | 0201g/0301g | 0303g/0602 | 1501g/5001 | 0401/0701 | 0202/0302 | 0201/0301 |
| CS5 | 0301g/3301 | 0702g/0802 | 0702g/1402 | 0301/1101 | 0201/0301 | 0501/0505 |
| CS6 | 0201g/0301g | 0602/0701g | 5701/5801g | 0701/0804 | 0303/0402 | 0201/0401 |
| NA12878 | 1101/0101 | 0102/0701 | 5601/0801 | 0101/0301 | 0501/0201 | 0101/0501 |

---

**Supplementary Figure 1.**
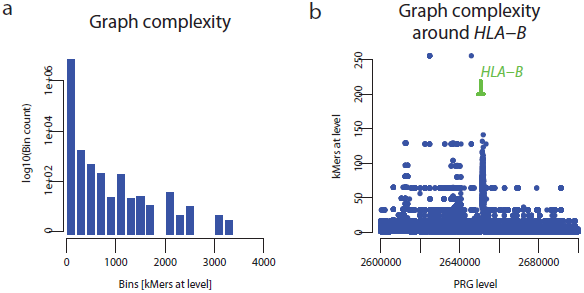
Complexity of the MHC population reference graph. **a.** Histogram showing the distribution of the number of kmers present in the PRG across sites within the extended MHC region. **b.** Spatial plot of graph complexity around *HLA-B* demonstrating peaks in complexity around classical HLA loci.

**Supplementary Figure 2.**
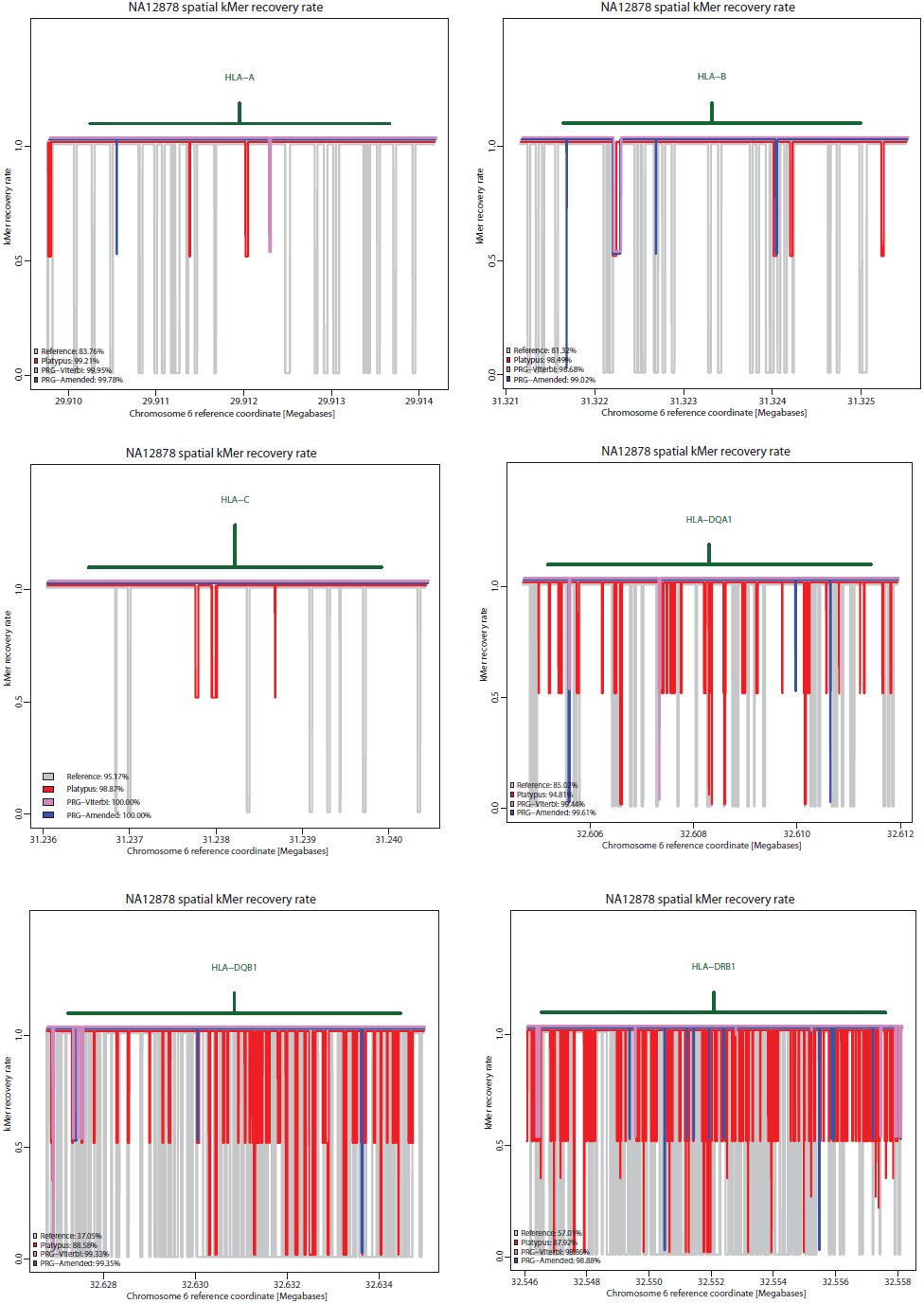
Kmer recovery within the classical HLA loci *HLA-A*, *-B*, *-C*, *-DQA1*, *-DQB1* and *-DRB1*. Each panel shows the fraction of kmers recovered at single nucleotide resolution from chromotypes inferred by the four methods using the high coverage data from NA12878. The average over the locus is also shown.

**Supplementary Table 1.** The types and counts of different types of input data used to construct the PRG for the human xMHC.

| Input data type |  | Number |
| --- | --- | --- |
| Scaffold haplotypes |  | 8 |
| SNPs |  | 87640 |
| Classical HLA alleles |  | 6602 |
| (genomic sequence) | HLAA | 1729 |
|  | HLAC | 1291 |
|  | HLAB | 2327 |
|  | HLADRB | 1051 |
|  | HLADQA | 46 |
|  | HLADQB | 158 |

**Supplementary Table 2.** Simulation-based assessment of PRG accuracy

| Category (per position) |  | Alleles evaluated | Alleles correct | Positions evaluated | Positions with 0/2 alleles correct | Positions with 1/2 alleles correct | Positions with 2/2 alleles correct |
| --- | --- | --- | --- | --- | --- | --- | --- |
| Graph structure constraints | True paths constraints |  |  |  |  |  |  |
| * | * | 199,937,840 | 99.86% | 99,968,920 | 0.03% | 0.23% | 99.74% |
| * | heterozygous | 24,051,922 | 99.26% | 12,025,961 | 0.00% | 1.49% | 98.51% |
| Clean SNP positions | * | 3,246,880 | 99.83% | 1,623,440 | 0.03% | 0.28% | 99.69% |
| Clean SNP positions | heterozygous | 1,560,760 | 99.83% | 780,380 | 0.00% | 0.33% | 99.67% |
| SV and INDEL positions | * | 52,485,120 | 99.46% | 26,242,560 | 0.11% | 0.86% | 99.03% |
| SV and INDEL positions | heterozygous | 22,473,412 | 99.22% | 11,236,706 | 0.00% | 1.57% | 98.43% |
| SV and INDEL positions | 1 "gap" genotype, heterozygous | 22,198,112 | 99.23% | 11,099,056 | 0.00% | 1.54% | 98.46% |
| SV and INDEL positions | 2 "gap" genotypes | 18,029,904 | 99.63% | 9,014,952 | 0.18% | 0.38% | 99.44% |
| SV and INDEL positions | 1 "gap" genotype, INDEL length 1 | 58,396 | 98.56% | 29,198 | 0.21% | 2.45% | 97.34% |
| SV and INDEL positions | 1 "gap" genotype, SV length 5 | 22,370 | 98.29% | 11,185 | 0.15% | 3.11% | 96.74% |
| SV and INDEL positions | 1 "gap" genotype, heterozygous, within 10-19 SV region | 89,044 | 95.76% | 44,522 | 0.00% | 8.49% | 91.51% |
| SV and INDEL positions | 1 "gap" genotype, heterozygous, within 100-149 SV region | 260,550 | 96.23% | 130,275 | 0.00% | 7.53% | 92.46% |
| SV and INDEL positions | 1 "gap" genotype, heterozygous, within 1000-1999 SV region | 1,432,310 | 99.37% | 716,155 | 0.00% | 1.26% | 98.74% |
| SV and INDEL positions | 1 "gap" genotype, heterozygous, within 60,000 - 60,999 SV region | 1,029,088 | 99.97% | 514,544 | 0.00% | 0.06% | 99.94% |
| SV and INDEL positions | 1 "gap" genotype, heterozygous, within 124,000 - 124,999 SV region | 347,462 | 99.88% | 173,731 | 0.00% | 0.23% | 99.77% |
| SV and INDEL positions | 1 "gap" genotype, heterozygous, within 10,000 - 10,999 SV region | 284,798 | 100.00% | 142,399 | 0.00% | 0.00% | 100.00% |

This table shows concordance between simulated true and inferred chromatypes (inferred from the reads simulated from the simulated true chromatypes; read length = 85bp, 30x diploid coverage; as the main effect of read errors is a slight reduction of kMer coverage, simulation was carried out without read error), stratified by the local properties of the chromatypes as well as of the graph at the evaluated positions.
Each position is evaluated as a diploid genotype consisting of two one-character alleles.
“Graph structure constraints”: Properties of the utilized PRG at the level of the evaluated position. “\*” means no constraints, “Clean SNP positions” are levels containing only edges labeled with “A”, “C”, “G” or “T”, “SV and INDEL positions” denote levels at which at least one edge is labeled with the gap character “\_”. “True path constraints” refers to the properties of the simulated (true) chromotype at the evaluated position. “\*” means no constraints; “heterozygous” means that the chromotype has two edges with different labels; “1 gap genotypes” means that the chromotype carries one gap label at the specified position and 1 non-gap label; “2 gap genotypes” means that the chromotype carries two gap symbols at the specified position; “within x – y SV region” means that it is possible to walk along the chromotype from the evaluated position a positions to the left and b positions to the right until hitting a position at which the chromotype does not carry a gap symbol, and $x < (a + b) \leq y$ .
“Alleles evaluated” is 2 x “Positions evaluated”. The remaining column headers are self-explanatory.

**Supplementary Table 3.**
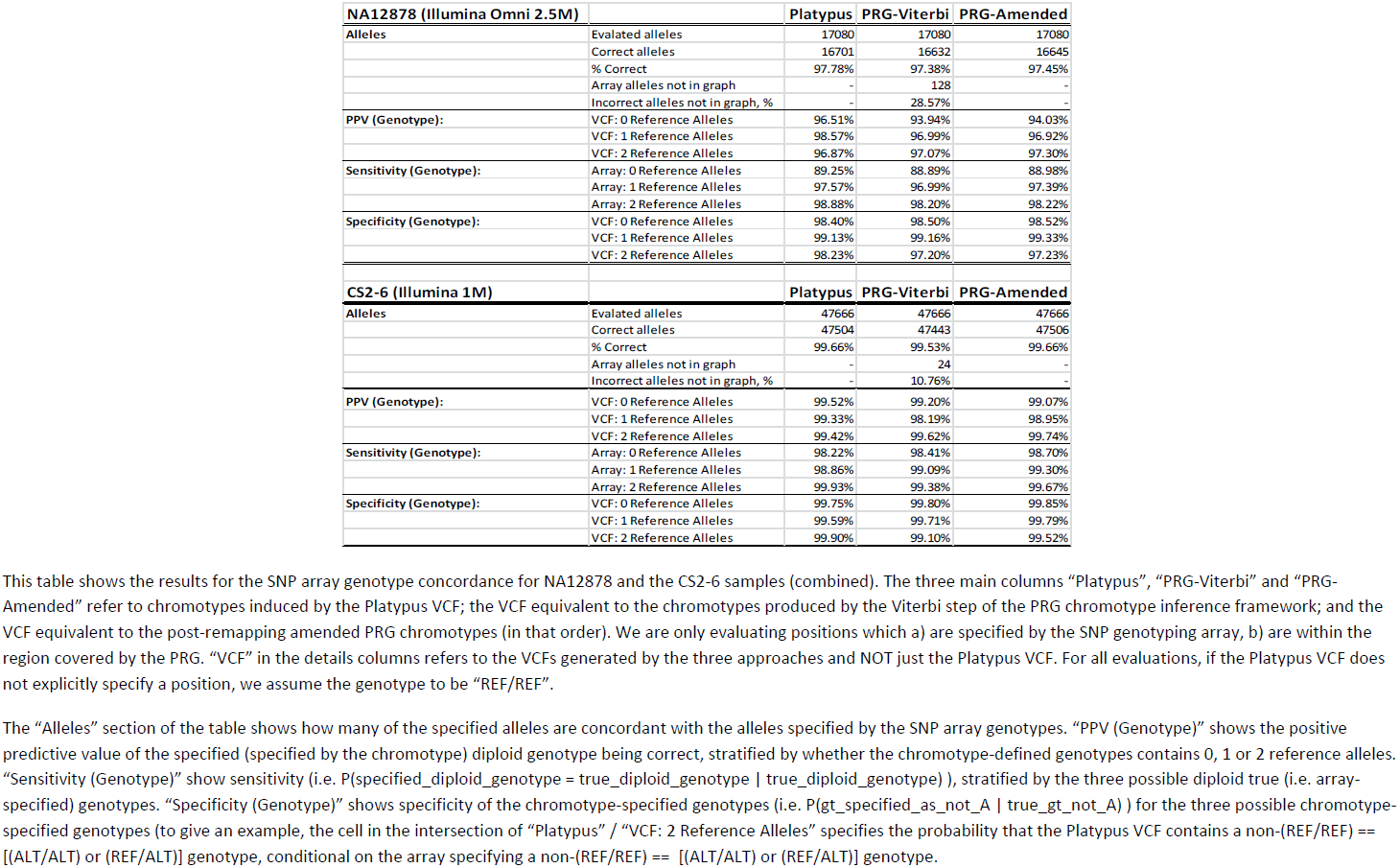
Accuracy of approaches assessed from SNP array data

| **NA12878 (Illumina Omni 2.5M)** |  | **Platypus** | **PRG-Viterbi** | **PRG-Amended** |
| --- | --- | --- | --- | --- |
| **Alleles** | Evalated alleles | 17080 | 17080 | 17080 |
|  | Correct alleles | 16701 | 16632 | 16645 |
|  | % Correct | 97.78% | 97.38% | 97.45% |
|  | Array alleles not in graph | - | 128 | - |
|  | Incorrect alleles not in graph, % | - | 28.57% | - |
| **PPV (Genotype):** | VCF: 0 Reference Alleles | 96.51% | 93.94% | 94.03% |
|  | VCF: 1 Reference Alleles | 98.57% | 96.99% | 96.92% |
|  | VCF: 2 Reference Alleles | 96.87% | 97.07% | 97.30% |
| **Sensitivity (Genotype):** | Array: 0 Reference Alleles | 89.25% | 88.89% | 88.98% |
|  | Array: 1 Reference Alleles | 97.57% | 96.99% | 97.39% |
|  | Array: 2 Reference Alleles | 98.88% | 98.20% | 98.22% |
| **Specificity (Genotype):** | VCF: 0 Reference Alleles | 98.40% | 98.50% | 98.52% |
|  | VCF: 1 Reference Alleles | 99.13% | 99.16% | 99.33% |
|  | VCF: 2 Reference Alleles | 98.23% | 97.20% | 97.23% |
| **CS2-6 (Illumina 1M)** |  | **Platypus** | **PRG-Viterbi** | **PRG-Amended** |
| **Alleles** | Evalated alleles | 47666 | 47666 | 47666 |
|  | Correct alleles | 47504 | 47443 | 47506 |
|  | % Correct | 99.66% | 99.53% | 99.66% |
|  | Array alleles not in graph | - | 24 | - |
|  | Incorrect alleles not in graph, % | - | 10.76% | - |
| **PPV (Genotype):** | VCF: 0 Reference Alleles | 99.52% | 99.20% | 99.07% |
|  | VCF: 1 Reference Alleles | 99.33% | 98.19% | 98.95% |
|  | VCF: 2 Reference Alleles | 99.42% | 99.62% | 99.74% |
| **Sensitivity (Genotype):** | Array: 0 Reference Alleles | 98.22% | 98.41% | 98.70% |
|  | Array: 1 Reference Alleles | 98.86% | 99.09% | 99.30% |
|  | Array: 2 Reference Alleles | 99.93% | 99.38% | 99.67% |
| **Specificity (Genotype):** | VCF: 0 Reference Alleles | 99.75% | 99.80% | 99.85% |
|  | VCF: 1 Reference Alleles | 99.59% | 99.71% | 99.79% |
|  | VCF: 2 Reference Alleles | 99.90% | 99.10% | 99.52% |

This table shows the results for the SNP array genotype concordance for NA12878 and the CS2-6 samples (combined). The three main columns “Platypus”, “PRG-Viterbi” and “PRG-Amended” refer to chromotypes induced by the Platypus VCF; the VCF equivalent to the chromotypes produced by the Viterbi step of the PRG chromotype inference framework; and the VCF equivalent to the post-remapping amended PRG chromotypes (in that order). We are only evaluating positions which a) are specified by the SNP genotyping array, b) are within the region covered by the PRG. “VCF” in the details columns refers to the VCFs generated by the three approaches and NOT just the Platypus VCF. For all evaluations, if the Platypus VCF does not explicitly specify a position, we assume the genotype to be “REF/REF”.
The “Alleles” section of the table shows how many of the specified alleles are concordant with the alleles specified by the SNP array genotypes. “PPV (Genotype)” shows the positive predictive value of the specified (specified by the chromotype) diploid genotype being correct, stratified by whether the chromotype-defined genotypes contains 0, 1 or 2 reference alleles. “Sensitivity (Genotype)” show sensitivity (i.e. $P(\text{specified\_diploid\_genotype} = \text{true\_diploid\_genotype} \mid \text{true\_diploid\_genotype})$ ), stratified by the three possible diploid true (i.e. array-specified) genotypes. “Specificity (Genotype)” shows specificity of the chromotype-specified genotypes (i.e. $P(\text{gt\_specified\_as\_not\_A} \mid \text{true\_gt\_not\_A})$ ) for the three possible chromotype-specified genotypes (to give an example, the cell in the intersection of “Platypus” / “VCF: 2 Reference Alleles” specifies the probability that the Platypus VCF contains a non-(REF/REF) == [(ALT/ALT) or (REF/ALT)] genotype, conditional on the array specifying a non-(REF/REF) == [(ALT/ALT) or (REF/ALT)] genotype.

**Supplementary Table 4.**
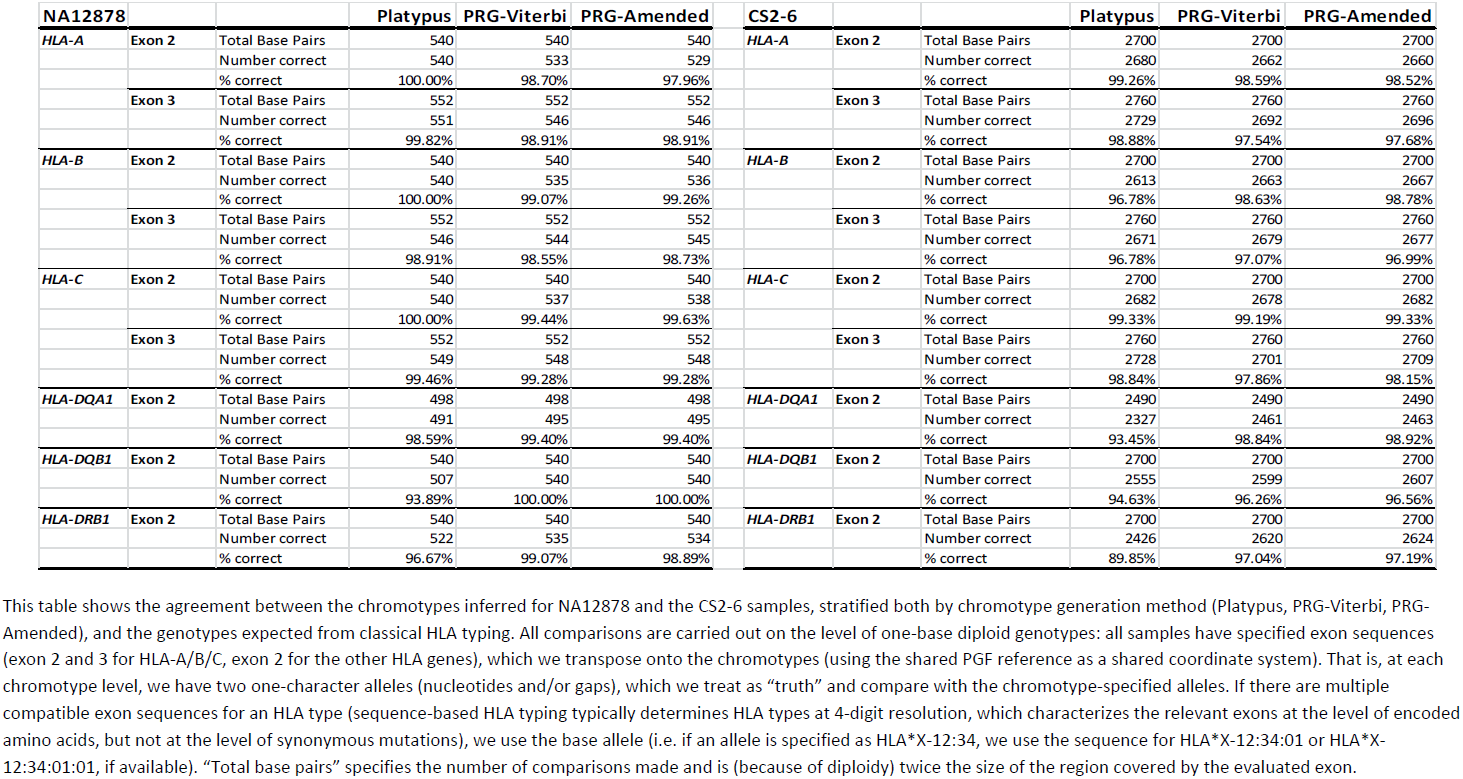
Accuracy of approaches assessed from classical HLA data

**Supplementary Table 5.** Accuracy of approaches estimated from kmer recovery

|  |  |  |  |  |  |
| --- | --- | --- | --- | --- | --- |
| **NA12878** |  |  |  |  |  |
|  | **Approach** | **Total # kMers** | **Invalid kMers** | **kMers present** | **Unweighted optimality** |
|  | PGF reference | 4518629 | 0 | 4301668 | 95.20% |
|  | Platypus VCF | 4938471 | 0 | 4880820 | 98.83% |
|  | PRG-Viterbi | 4984797 | 118 | 4953174 | 99.37% |
|  | PRG-Remapped | 4971054 | 116 | 4942609 | 99.43% |
| **CS2-6** |  |  |  |  |  |
| **CS2** |  |  |  |  |  |
|  | **Method** | **# kMers** | **# kMers invalid** | **# kMers present** | **Unweighted optimality** |
|  | PGF reference | 4518629 | 0 | 4255288 | 94.17% |
|  | Platypus VCF | 4950328 | 0 | 4860657 | 98.19% |
|  | PRG-Viterbi | 5073842 | 647 | 5042634 | 99.38% |
|  | PRG-Remapped | 5051678 | 647 | 5022085 | 99.41% |
| **CS3** |  |  |  |  |  |
|  | **Method** | **# kMers** | **# kMers invalid** | **# kMers present** | **Unweighted optimality** |
|  | PGF reference | 4518629 | 0 | 4248008 | 94.01% |
|  | Platypus VCF | 4889852 | 0 | 4801145 | 98.19% |
|  | PRG-Viterbi | 5015592 | 824 | 4992734 | 99.54% |
|  | PRG-Remapped | 4989059 | 824 | 4966451 | 99.55% |
| **CS4** |  |  |  |  |  |
|  | **Method** | **# kMers** | **# kMers invalid** | **# kMers present** | **Unweighted optimality** |
|  | PGF reference | 4518629 | 0 | 4282989 | 94.79% |
|  | Platypus VCF | 4830916 | 0 | 4729207 | 97.89% |
|  | PRG-Viterbi | 4931881 | 575 | 4893711 | 99.23% |
|  | PRG-Remapped | 4892548 | 575 | 4860145 | 99.34% |
| **CS5** |  |  |  |  |  |
|  | **Method** | **# kMers** | **# kMers invalid** | **# kMers present** | **Unweighted optimality** |
|  | PGF reference | 4518629 | 0 | 4344265 | 96.14% |
|  | Platypus VCF | 4881754 | 0 | 4794764 | 98.22% |
|  | PRG-Viterbi | 4860787 | 627 | 4839312 | 99.56% |
|  | PRG-Remapped | 4837182 | 627 | 4817300 | 99.59% |
| **CS6** |  |  |  |  |  |
|  | **Method** | **# kMers** | **# kMers invalid** | **# kMers present** | **Unweighted optimality** |
|  | PGF reference | 4518629 | 0 | 4273214 | 94.57% |
|  | Platypus VCF | 4913174 | 0 | 4818055 | 98.06% |
|  | PRG-Viterbi | 5015081 | 518 | 4971576 | 99.13% |
|  | PRG-Remapped | 4990040 | 518 | 4951327 | 99.22% |
| **CS2-6 comb.** |  |  |  |  |  |
|  | **Method** | **# kMers** | **# kMers invalid** | **# kMers present** | **Unweighted optimality** |
|  | PGF reference | 22,593,145 | 0 | 21,403,764 | 94.74% |
|  | Platypus VCF | 24,466,024 | 0 | 24,003,828 | 98.11% |
|  | PRG-Viterbi | 24,897,183 | 3,191 | 24,739,967 | 99.37% |

This table shows the results for kMer evaluation for the four chromatypes (PRG-Viterbi and PRG-Amended, the chromotype constructed from the VCF covering the PRG-covered region, and the [non-branching] chromotype constructed from the PGF reference for the region covered by the PRG).
For any two positions along a chromotype where phase is lost (i.e. at positions where one node in the chromotype has two outgoing edges) within 31 bases distance, we need to resolve phase before applying the evaluation algorithm. We use a simple greedy algorithm which keeps track of the possible haplotype pairs induced by chromotype structure, until either the chromotype has reached a homozygous (i.e. one node, one edge at each level) stretch of 31 defined base pairs length (i.e. at least 31 non-gap characters) or the number of possible haplotype pairs has reached a threshold (see Supplementary Methods). At this point, we count how many of the kMers induced by each haplotype pair are present in the sample, and fix the pair with the highest number of kMers present in the sample. This process does not affect the “PGF reference” and the “PRG-Viterbi” chromatypes (because kMerification ensures that there is no loss of phase within the specified kMer size).
As we have disentangled all positions with the potential to locally disrupt haplotypes, we can now count how often each kMer appears along a chromotype. “Unweighted optimality” is the percentage of the kMers present also found in the sample sequencing data.
“#kMers invalid” counts how many of the specified kMers are invalid, i.e. containing a non-nucleotide “\*” symbol (the PRG construction algorithm tries to remove edges labeled with the “\*” symbol, but not always successfully). Invalid kMers are counted as “not present”.

**Supplementary Table 6.** Alignment metrics for Moleculo contig alignment to chromotypes

| Method | TotalAlignmentLength | AverageAlignmentSequenceCertainty | AveragekMerUniqueness |
| --- | --- | --- | --- |
| Reference | 566,390,766 | 0.99 | 0.85 |
| VCF | 365,509,324 | 0.97 | 0.93 |
| PRG-Viterbi | 294,127,027 | 0.99 | 0.86 |
| PRG-Amended | 272,355,664 | 0.97 | 0.94 |
| Method | graphIntrinsicGap_sequenceGap | graphIntrinsicGap_sequenceCharacter | Matches |
| Reference | 0 | 0 | 114,518,701 |
| VCF | 555,158 | 109 | 115,059,507 |
| PRG-Viterbi | 30,268,853 | 844 | 115,688,453 |
| PRG-Amended | 25,871,037 | 1,218 | 115,519,131 |
| Method | Mismatches | graphNovelGap | graphNonGap_sequenceGap |
| Reference | 540,803 | 1,101,559 | 450,229,703 |
| VCF | 84,216 | 1,017,231 | 248,793,103 |
| PRG-Viterbi | 91,288 | 380,478 | 147,697,111 |
| PRG-Amended | 53,862 | 586,852 | 130,323,564 |

This table shows the results from the alignment of the 29429 Molecuro contigs (median contig length 3165bp) to chromotypes from four sources. The numbers displayed are cumulative, i.e. **sums over the individual values for all contigs**.
“TotalAlignmentLength” specifies the total length of the contig alignments, measured as columns in the alignments (the alignment algorithm optimizes for global alignment, but truncates the alignments at the positions of the last aligned contig characters).
“AverageAlignmentSequenceCertainty” specifies how certain the algorithm is in its assignment of contig characters to chromotype levels (using a randomized sampling procedure, see Supplementary text).
“AveragekMerUniqueness” specifies the percentage of kMers in the contig that are double-unique, i.e. unique in both contig and chromotype that the contig is aligned to. Double-unique kMers are used as initial anchors in the alignment procedure.
“graphIntrinsicGap\_sequenceGap” specifies how many columns in the alignment specify a graph gap symbol (i.e. a gap present in the PRG / VCF) in the chromotype row and gap symbol in the contig row.
“graphIntrinsicGap\_sequenceCharacter” specifies how many columns in the alignment specify a graph gap symbol (i.e. a gap present in the PRG / VCF) in the chromotype row and a non-gap symbol in the contig row.
“Matches” counts how many positions in the alignments specify the same non-gap symbol for chromotype and contig.
“Mismatches” counts how many positions in the alignments specify a pair (a, b) of symbols for chromotype and contig, with a != b and neither a nor b equal to a gap character.
“graphNovelGap” counts how many positions in the alignments specify a non-gap symbol in the contig row and a novel gap in the chromotype (novel gap := gap symbol not coming from the PRG / VCF). These positions are indicative of insertions in the contig relative to the chromotype (or contig misassembly or alignment problems).

## References

1. Li, H. & Durbin, R. Fast and accurate long-read alignment with Burrows-Wheeler transform. Bioinformatics 26, 589–95 (2010).

2. Li, H., Ruan, J. & Durbin, R. Mapping short DNA sequencing reads and calling variants using mapping quality scores. Genome Res 18, 1851–8 (2008).

3. Lunter, G. & Goodson, M. Stampy: a statistical algorithm for sensitive and fast mapping of Illumina sequence reads. Genome Res 21, 936–9 (2011).

4. Horton, R. et al. Variation analysis and gene annotation of eight MHC haplotypes: the MHC Haplotype Project. Immunogenetics 60, 1–18 (2008).

5. Jiang, W. et al. Copy number variation leads to considerable diversity for B but not A haplotypes of the human KIR genes encoding NK cell receptors. Genome Res 22, 1845–54 (2012).

6. Trask, B.J. et al. Large multi-chromosomal duplications encompass many members of the olfactory receptor gene family in the human genome. Hum Mol Genet 7, 2007–20 (1998).

7. Steinberg, K.M. et al. Structural diversity and African origin of the 17q21.31 inversion polymorphism. Nat Genet 44, 872–80 (2012).

8. Boettger, L.M., Handsaker, R.E., Zody, M.C. & McCarroll, S.A. Structural haplotypes and recent evolution of the human 17q21.31 region. Nat Genet 44, 881–5 (2012).

9. Stefansson, H. et al. A common inversion under selection in Europeans. Nat Genet 37, 129–37 (2005).

10. Lupski, J.R. & Stankiewicz, P. Genomic disorders: molecular mechanisms for rearrangements and conveyed phenotypes. PLoS Genet 1, e49 (2005).

11. The Genome Reference Consortium. http://www.ncbi.nlm.nih.gov/assembly/GCF_000001405.26/. (2014).

12. The 1000 Genomes Project Consortium. A map of human genome variation from population-scale sequencing. Nature 467, 1061–73 (2010).

13. The International HapMap Consortium. A haplotype map of the human genome. Nature 437, 1299–320 (2005).

14. The 1000 Genomes Project Consortium. An integrated map of genetic variation from 1,092 human genomes. Nature 491, 56–65 (2012).

15. Rimmer, A. et al. Integrating mapping, assembly and haplotype-based approaches for calling variants in clinical sequencing applications. Nat Genet (In press).

16. Garrison, E.P. & Marth, G. Haplotype-based variant detection from short-read sequencing. (http://arxiv.org/abs/1207.3907, 2012).

17. Kural, D. & Garrison, E.P. (https://github.com/ekg/glia).

18. Huang, L., Popic, V. & Batzoglou, S. Short read alignment with populations of genomes. Bioinformatics 29, i361–70 (2013).

19. Schneeberger, K. et al. Simultaneous alignment of short reads against multiple genomes. Genome Biol 10, R98 (2009).

20. Sirén, J., Valimäki, N. & Mäkinen, V. Indexing finite language representation of population genotypes. Proc. WABI 6833, 270–281 (2011).

21. Paten, B., Novak, A. & Haussler, D. Mapping to a reference genome structure. (http://arxiv.org/abs/1404.5010, 2014).

22. Iqbal, Z., Caccamo, M., Turner, I., Flicek, P. & McVean, G. De novo assembly and genotyping of variants using colored de Bruijn graphs. Nat Genet 44, 226–32 (2012).

23. Katoh, K. & Frith, M.C. Adding unaligned sequences into an existing alignment using MAFFT and LAST. Bioinformatics 28, 3144–6 (2012).

24. Bradley, R.K. et al. Fast statistical alignment. PLoS Comput Biol 5, e1000392 (2009).

25. Lefranc, M.P. et al. IMGT, the international ImMunoGeneTics information system. Nucleic acids research 37, D1006–12 (2009).

26. Talwalkar, A. et al. SMaSH: A Benchmarking Toolkit for Human Genome Variant Calling. (http://arxiv.org/abs/1310.8420, 2014).

27. Simpson, J.T. & Durbin, R. Efficient de novo assembly of large genomes using compressed data structures. Genome Res 22, 549–56 (2012).

28. Li, Y., Willer, C.J., Ding, J., Scheet, P. & Abecasis, G.R. MaCH: using sequence and genotype data to estimate haplotypes and unobserved genotypes. Genetic Epidemiology 34, 816–34 (2010).

29. Dilthey, A. et al. Multi-population classical HLA type imputation. PLoS Comput Biol 9, e1002877 (2013).

30. Howie, B., Marchini, J. & Stephens, M. Genotype imputation with thousands of genomes. G3 1, 457–70 (2011).

31. Browning, S.R. & Browning, B.L. Rapid and accurate haplotype phasing and missing-data inference for whole-genome association studies by use of localized haplotype clustering. Am J Hum Genet 81, 1084–97 (2007).

32. Li, Y., Sidore, C., Kang, H.M., Boehnke, M. & Abecasis, G.R. Low-coverage sequencing: implications for design of complex trait association studies. Genome Res 21, 940–51 (2011).

33. Spraggs, C.F., Parham, L.R., Hunt, C.M. & Dollery, C.T. Lapatinib-induced liver injury characterized by class II HLA and Gilbert’s syndrome genotypes. Clin Pharmacol Ther 91, 647–52 (2012).

34. Lunter, G. & Goodson, M. Stampy: A statistical algorithm for sensitive and fast mapping of Illumina sequence reads. Genome Res (2010).

## References

1. Dilthey, A., et al., Multi-Population Classical HLA Type Imputation. PLoS Comput Biol, 2013. 9(2): p. e1002877.

2. Browning, S.R. and B.L. Browning, Rapid and accurate haplotype phasing and missing-data inference for whole-genome association studies by use of localized haplotype clustering. Am J Hum Genet, 2007. 81(5): p. 1084–97.

3. Rabiner, L.R., A tutorial on hidden Markov models and selected applications in speech recognition. Proceedings of the Ieee, 1989. 77(2): p. 257–286.

4. Needleman, S.B. and C.D. Wunsch, A general method applicable to the search for similarities in the amino acid sequence of two proteins. J Mol Biol, 1970. 48(3): p. 443–53.

5. Li, H. and R. Durbin, Fast and accurate long-read alignment with Burrows-Wheeler transform. Bioinformatics, 2010. 26(5): p. 589–95.

6. Li, H., et al., The Sequence Alignment/Map format and SAMtools. Bioinformatics, 2009. 25(16): p. 2078–9.

7. Sankoff, D., Matching sequences under deletion-insertion constraints. Proc Natl Acad Sci U S A, 1972. 69(1): p. 4–6.

8. Horton, R., et al., Variation analysis and gene annotation of eight MHC haplotypes: the MHC Haplotype Project. Immunogenetics, 2008. 60(1): p. 1–18.

9. Bradley, R.K., et al., Fast statistical alignment. PLoS Comput Biol, 2009. 5(5): p. e1000392.

10. Katoh, K. and M.C. Frith, Adding unaligned sequences into an existing alignment using MAFFT and LAST. Bioinformatics, 2012. 28(23): p. 3144–6.

11. Genomes Project, C., et al., An integrated map of genetic variation from 1,092 human genomes. Nature, 2012. 491(7422): p. 56–65.

12. Holdsworth, R., et al., The HLA dictionary 2008: a summary of HLA-A, -B, -C, -DRB1/3/4/5, and -DQB1 alleles and their association with serologically defined HLA-A, -B, -C, -DR, and -DQ antigens. Tissue Antigens, 2009. 73(2): p. 95–170.

13. Flicek, P., et al., Ensembl 2013. Nucleic Acids Res, 2013. 41(Database issue): p. D48–55.

